# *De novo* design of ACE2 protein decoys to neutralize SARS-CoV-2

**DOI:** 10.1101/2020.08.03.231340

**Authors:** Thomas W. Linsky, Renan Vergara, Nuria Codina, Jorgen W. Nelson, Matthew J. Walker, Wen Su, Tien-Ying Hsiang, Katharina Esser-Nobis, Kevin Yu, Yixuan J. Hou, Tanu Priya, Masaya Mitsumoto, Avery Pong, Uland Y. Lau, Marsha L. Mason, Jerry Chen, Alex Chen, Tania Berrocal, Hong Peng, Nicole S. Clairmont, Javier Castellanos, Yu-Ru Lin, Anna Josephson-Day, Ralph Baric, Carl D. Walkey, Ryan Swanson, Michael Gale, Luis M. Blancas-Mejia, Hui-Ling Yen, Daniel-Adriano Silva

## Abstract

There is an urgent need for the ability to rapidly develop effective countermeasures for emerging biological threats, such as the severe acute respiratory syndrome coronavirus 2 (SARS-CoV-2) that causes the ongoing coronavirus disease 2019 (COVID-19) pandemic. We have developed a generalized computational design strategy to rapidly engineer *de novo* proteins that precisely recapitulate the protein surface targeted by biological agents, like viruses, to gain entry into cells. The designed proteins act as decoys that block cellular entry and aim to be resilient to viral mutational escape. Using our novel platform, in less than ten weeks, we engineered, validated, and optimized *de novo* protein decoys of human angiotensin-converting enzyme 2 (hACE2), the membrane-associated protein that SARS-CoV-2 exploits to infect cells. Our optimized designs are hyperstable de novo proteins (∼18-37 kDa), have high affinity for the SARS-CoV-2 receptor binding domain (RBD) and can potently inhibit the virus infection and replication in vitro. Future refinements to our strategy can enable the rapid development of other therapeutic *de novo* protein decoys, not limited to neutralizing viruses, but to combat any agent that explicitly interacts with cell surface proteins to cause disease.

## Main Text

Since its emergence in December of 2019, SARS-CoV-2 has caused millions of cases of COVID-19 and become a global pandemic. The need for effective strategies to prevent and treat the disease remains unfulfilled and urgent (*1*). There are multiple ongoing efforts to develop prophylactics and therapeutics using several approaches (*2*) such as vaccination (*3*), protein engineering (*1*) and small molecule drug discovery (*4*). The high mutational rate of positive sense single-strand RNA (+ssRNA) viruses (*5–7*) can lead to viral escape (*8*), which could rapidly compromise the efficacy of many treatments under development, possibly leading to recurrent outbreaks of SARS-CoV-2 with globally catastrophic consequences. Mutations have already spontaneously occurred in the spike (S) protein of SARS-CoV-2 (*9*), and some of them are already suspected to increase the infectivity of the virus (*10*)). Deep mutational scanning studies have further shown that there are mutations that can enable the virus to escape known neutralizing antibodies, as well as mutations that can increase its binding affinity for hACE2 (*11, 12*). A pressing need exists to develop novel therapeutics that can be more resistant to viral mutation and immunologic escape by SARS-CoV-2. Traditional approaches to combat viruses (e.g. monoclonal antibodies) ultimately rely on molecules interacting with viruses in a way that is fundamentally different than how the virus engages with its cellular targets (*13, 14*). Viruses can exploit such structural discrepancy to evade neutralization, changing the shape of their proteins to prevent recognition by the neutralizing agents while preserving their viral function. To address these challenges, we have developed a general protein design strategy that enables the rapid and accurate computational design of *de novo* proteins that neutralize a virus by disabling its mechanism of cellular entry. By mimicking the exact protein surface that the virus targets on its first contact with the cell’s membrane, our *de novo* designer proteins can outcompete the viral interaction and act as neutralizing agents. Since a de novo protein decoy mirrors the natural viral cell’s protein target, mutations that affect (e.g. weaken) the interaction of the virus with the *de novo* decoy are likely to have a similar effect on its interaction with the cell, automatically conferring the designed decoys with resiliency to mutational viral escape.

SARS-CoV-2 invades host cells in a two-step process (*15–17*). The first step occurs when the SARS-CoV-2 S protein attaches to the cell via binding to the angiotensin converting enzyme 2 (ACE2), triggering endocytosis and the second when it escapes the endosome via a protease-cleavage-mediated fusion peptide (*18*). The process is similar to the beta-coronaviruses HCoV-NL63 and SARS-CoV-1, which also target ACE2 for cellular entry (*19*). In principle, inhibiting the viral interaction with ACE2 should therefore prevent infection.In a span of ten weeks (figure 1J), we applied our design strategy to engineer, validate and optimize *de novo* protein decoys of human ACE2 (hACE2) to neutralize SARS-CoV-2 (figure S1). Our best optimized design, CTC-445.2d (ConquerTheCorona445.2duo), is a thermodynamically hyperstable, bi-valent, single-chain protein that exhibits low nanomolar affinity for the RBD of SARS-CoV-2 and is cross-reactive with SARS-CoV-1. As predicted, the strong and specific binding of CTC-445.2d to the RBD of SARS-CoV-2 translated into potent and specific *in vitro* viral neutralization. As expected, the effect SARS-CoV-2 RBD mutations in binding are closely mirrored between hACE2 and the de novo decoys.At a size of ∼160 a.a. and combining multiple α-helical and β-strand structures, CTC-445 and its optimized variants are among the most advanced examples of *de novo* therapeutic proteins reported to date.

**Figure 1.**
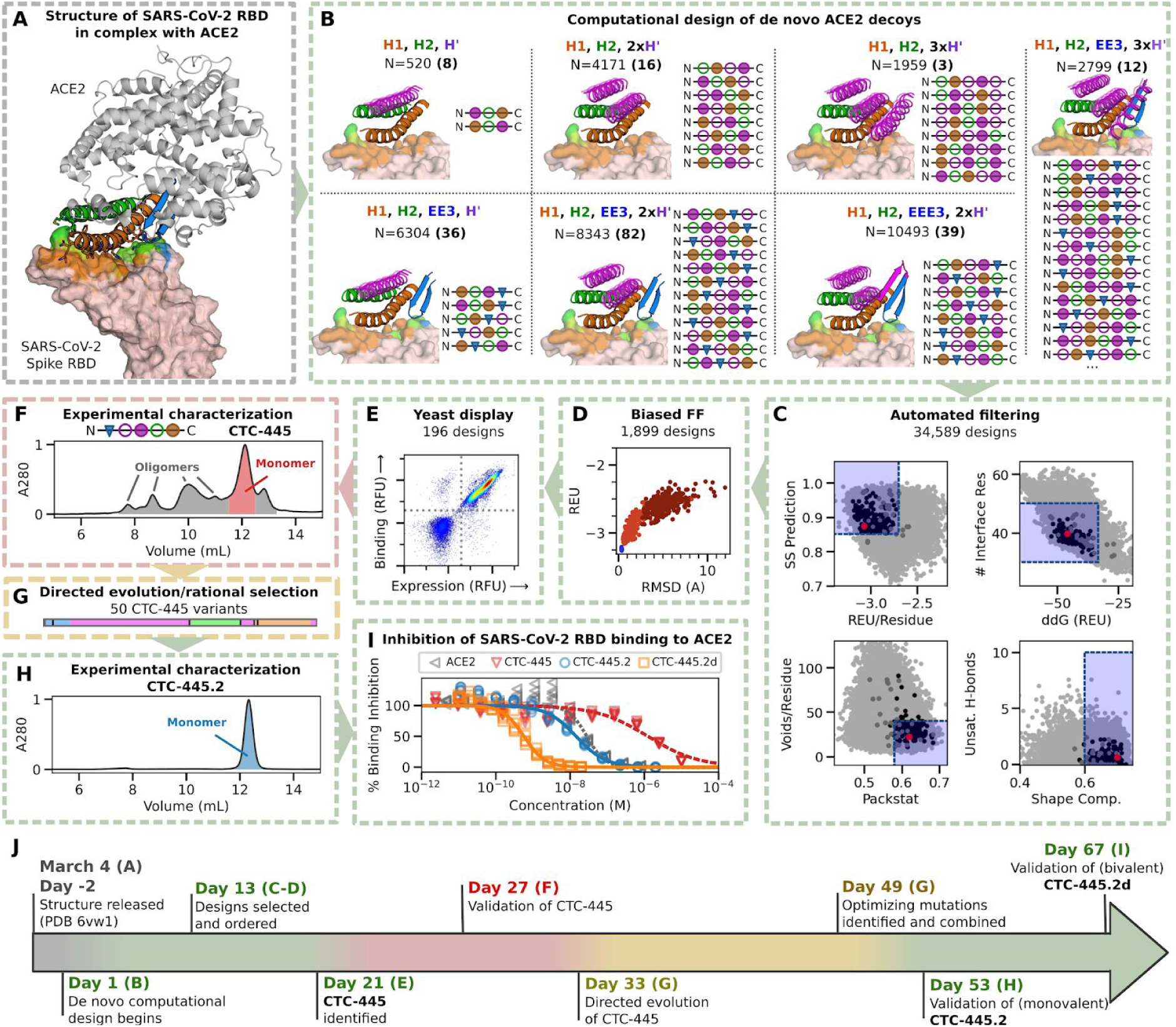
Design and characterization of *de novo* ACE2 decoys. **A)** ACE2 (gray) and its binding motifs (H1 19-52, orange; H2 55-84, green; EE3 346-360, blue) to SARS-CoV-2 RBD (pink). Three starting structures were simultaneously used as the target (see main text), 6VW1 is shown; **B)** Strategies for supporting the binding motifs. *De novo* secondary structure elements (magenta) were computationally generated to stabilize H1, H2, and EE3. Seven combinations of secondary structure elements were considered. Circles indicate *α*-helices, triangles indicate *β*-sheets; filled circles are helices oriented forward and empty circles are the opposite. We used Rosetta to generate fully connected backbones (“protein_mimic_designer” algorithm) and amino acid sequences predicted to fold into the target structure. In all cases, the binding interface of ACE2 with the SARS-CoV-2 RBD was preserved intact (see methods); **C)** Automatic computational filtering based on 8 metrics to select the best candidates. In addition, the RMSD of the binding motifs to ACE2 was also used as a quality check. The dots indicate the mean computational score for each design scored against the three target RBD structures. Designs selected for experimental testing are shown in black. Our best design, CTC-445, is shown in red. The blue boxes indicate the filtering thresholds (see methods); **D)** Designs that passed filtering were subjected to biased forward folding simulations (see methods), here shown for CTC-445, including the unsalted biased simulation (brown), the native-salted (orange), and relaxation (blue). **E)** The top 196 designs were selected for yeast display screening using a combination of Rosetta score per residue, ddG, and the folding simulations (see methods); The designs were individually assessed for specific binding to SARS-CoV-2 spike RBD (Fc fusion, 200 nM). The plot for CTC-445 is shown. **F)** CTC-445 was recombinantly expressed and purified by affinity chromatography (see methods). Analytical size exclusion chromatography (SEC) for CTC-445 revealed the presence of oligomeric species; **G)** CTC-445 was optimized by directed evolution and rational combination of the observed favorable mutations, leading to **H)** CTC-445.2 (SEC) which is mainly monomeric in solution and ∼1000x more potent to compete ACE2 than its parent (see “G”); we further optimized the potency of our molecule by generating a bivalent version named CTC-445.2d; **I)** Potency of designs to outcompete binding of SARS-CoV-2 to ACE2, as measured by competition ELISA assays using a constant concentration of 0.4 nM ACE2. **J)** Timeline of the *de novo* protein design and optimization pipeline. Timewise, a green color indicates phases that we believe were carried out optimally, red can potentially be avoided in future efforts, and yellow are phases that can potentially be expedited by using more advanced/automated methods for gene synthesis, cloning and high-throughput screening.

### Computational design and optimization of *de novo* ACE2 decoys to neutralize SARS-CoV-2

We started the design process of the de novo decoys by identifying the hACE2 structural motifs that form the protein surface that SARS-CoV-2 binds to gain entry into the cell. We based our effort on three publicly available structures of hACE2 in complex with the RBDs of SARS-CoV-1 (PDB: 6CS2) and SARS-CoV-2 (PDBs: 6VW1, 6M17) (*20–22*). By visual inspection, we identified four discontiguous hACE2 structural elements that form the target protein binding interface. We selected the major three of these elements to build our de novo decoys: two long alpha helices (H1 and H2) and a short beta hairpin (EE3) (figures 1A and S2). To generate molecules that are biologically inert for humans, our design strategy avoided incorporating elements of hACE2 that are known (or predicted) to be biologically active, such as its catalytic site and/or its interface with the cell. We built new disembodied *de novo* secondary structure elements tailored to support the target structural elements in a way that is both compatible with globular folding and that stabilizes the binding interface (figure 1B and methods), inspired by recent developments in the design of de novo structural elements (*23–26*). Then, in a spirit similar to the design of Neoleukin-2/15 (Neo-2/15) (*23, 27*), we used a combinatorial approach based on Rosetta’s “protein_mimic_designer” to generate multiple fully-connected protein topologies that contain all the desired structural and binding elements (*23*). The design of the protein decoys was constrained to fully preserve --intact up to each amino acid’s conformation-- the target binding interface (figure 1A-B, 2A-B), such as the designed de novo decoys can be more resilient to mutational escape by the virus. We then used Rosetta to generate de novo amino acid sequences predicted to fold into the target structures (figure 1B and methods), and evaluated the designs with an automatic filtering pipeline based on nine computational parameters, including predictions of smooth folding funnels into stable protein structures (figure 1C-D). The top ranking designs (see methods) were selected for experimental testing of binding to SARS-CoV-2 RBD (figure 1E).

We engineered ∼35,000 plausible computational models of *de novo* ACE2 decoys, and by using yeast display, we individually tested 196 of the top ranked designs for SARS-CoV-2 binding. Out-of-the-box, one of the designs, CTC-445, showed strong/nanomolar specific binding to the SARS-CoV-2 RBD (figure 1E, S3, and methods). Despite its high affinity, recombinantly expressed CTC-445 was a weak (µM range) competitor of hACE2 binding to SARS-CoV-2 RBD (figure 1I). A single round of directed evolution and rational combination of the most frequently observed (stabilizing and affinity-improving) mutations led to CTC-445.2, which contains five amino acid substitutions (figure 1G, S4, S5, and table S1, see methods). The improving mutations play a role in optimizing the stability of the protein’s native state, and none of them is at the binding interface (figures 2C-D, S4, and methods). CTC-445.2 is highly soluble, monodisperse (figure 1H and S7), thermodynamically hyperstable (figure 2C-D), a strong competitor of SARS-CoV-2 RBD binding to hACE2 (figure 1I), and a potent in vitro inhibitor of SARS-CoV-2 cell infection (figure 3). A simple domain duplication of CTC-445.2 (figure 2B) allowed us to further develop CTC-445.2d, which has similar favorable biochemical characteristics, but due to its bivalent nature exhibits a ∼100-fold improved SARS-CoV-2 neutralizing potency.

**Figure 2.**
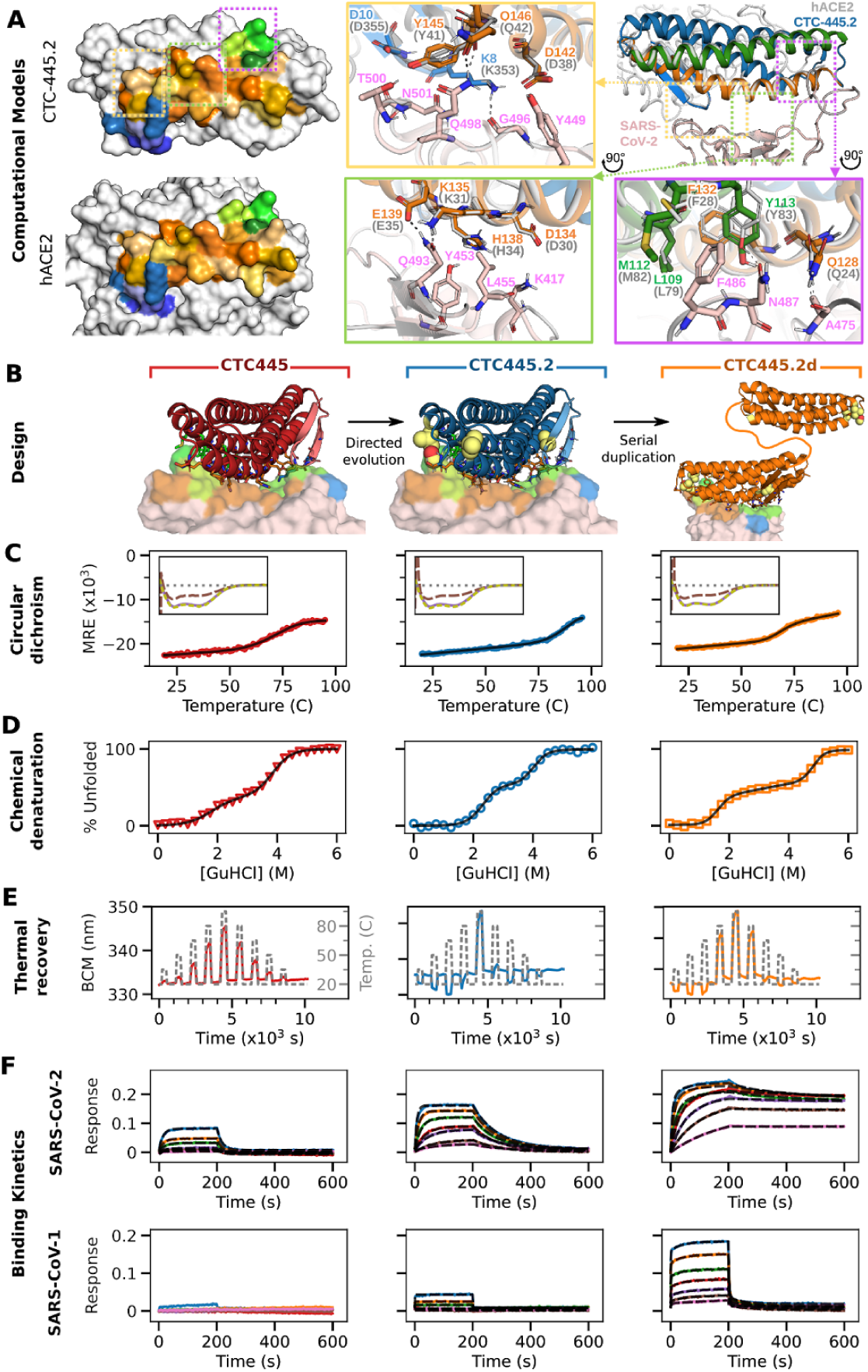
Binding, stability and structure of the *de novo* protein decoys CTC-445, CTC-445.2 and CTC-445.2d. **A)** Computational models of CTC-445.2. Left, comparison of the SARS-CoV-2 binding interface surface of CTC-445.2 (top) and hACE2 (bottom). CTC-445.2 is designed to present the same binding interface as the one that SARS-CoV-2 targets in hACE2, down to the level of individual interactions. Residues from the binding motifs: H1 are shown in shades of orange, residues from H2 in shades of green and residues from EE3 in shades of blue. The boxes show detailed structural comparison of the interfaces between CTC-445.2 and hACE2 with SARS-CoV-2 RBD. The relaxed complex of hACE2 with SARS-CoV-2 RBD (dark and light gray, respectively; PDB: 6M17) are aligned to the model of the relaxed compex of CTC-445.2 and SARS-CoV-2 RBD (pink). Hydrogen bond interactions are indicated by black dashed lines; **B)** Design models of CTC-445, CTC-445.2 and CTC-445.2d. CTC-445.2 contains 5 mutations that were guided by directed evolution experiments. CTC-445.2d is a bivalent variant composed of two CTC-445.2 subunits linked by a 17-mer flexible GS linker (sequence -GGGGSGGSGSGGSGGGS-); **C)** Circular dichroism of recombinantly expressed CTC-445 (red), CTC-445.2 (blue) and CTC-445.2d (orange). Thermally-induced meltings of the decoys were followed by its circular dichroism signal at 208 nm (heating rate 2°C/min). The inset shows far UV wavelength spectra at 20 °C (purple), after heating to ∼95 °C (brown) and after cooling the heated sample to 20 °C (green dashed). Complete ellipticity-spectra recovery (full reversibility) upon cooling was observed in all cases. Calculated T_m_ values for CTC-445, CTC-445.2 and 445.2d are 75.3 ± 0.2 °C, ≈93 °C and 71.7.± 0.2 °C; **D)** Chemical denaturation of CTC-445, CTC-445.2 and CTC-445.2d by incubation with guanidine hydrochloride (0-6 M). Calculated values of ΔG_ND_, ΔG_NI_ and ΔG_ID_ were -9.8 ± 0.8, -2.7 ± 1.1 and -7.1 ± 0.7 kcal mol^-1^ for CTC-445, -14.6 ± 2.0, -5.0 ± 2.8 and -9.6 ± 2.0 kcal mol^-1^ for CTC-445.2, and -19.0 ± 1.7, -4.4 ± 2.3 and -14.6 ± 1.6 kcal mol^-1^ for CTC-445.2d; **E)** Thermal recovery of CTC-445, CTC-445.2 and CTC-445.2d. Similar values of BCM were observed before and after repeated cycles of heating and cooling for the three proteins; **F)** Octet binding assays of CTC-445, CTC-445.2 (left) and CTC-445.2d (right) against immobilized SARS-CoV-2 RBD (top) and SARS-CoV-1 RBD (bottom). The RBD was immobilized to Anti-Penta-HIS (HIS1K, ForteBio) and incubated with varying concentrations of ACE2 decoys. Binding kinetics were monitored by dipping the biosensors in wells containing defined concentrations (4.7-300 nM for binding to SARS-CoV-2 and 30-2500 nM for binding to SARS-CoV-1) of CTC-445, CTC-445.2, CTC-445.2d (association) and then dipping the sensors back into baseline wells (dissociation). Data traces shown are a 2 s (10 data point) rolling average. Data for CTC-445 was fit to 1:1 model. For CTC-445 and CTC-445.2, the results were globally fit using a 1:1 binding model (K_d_=646 and 30 nM, k_on_=8.6 × 10^4^ and 3.9 × 10^5^ M^-1^s^-1^, k_off_ =5.5 × 10^−2^ and 1.2 × 10^−2^ s^-1^) and. For CTC-445.2d, the results were globally fit using a 2:1 binding model (K_d_ ≤7.0 nM, k_on_=3.0 × 10^5^ M^-1^s^-1^, k_off_≤2.0 × 10^−3^ s^-1^).

**Figure 3.**
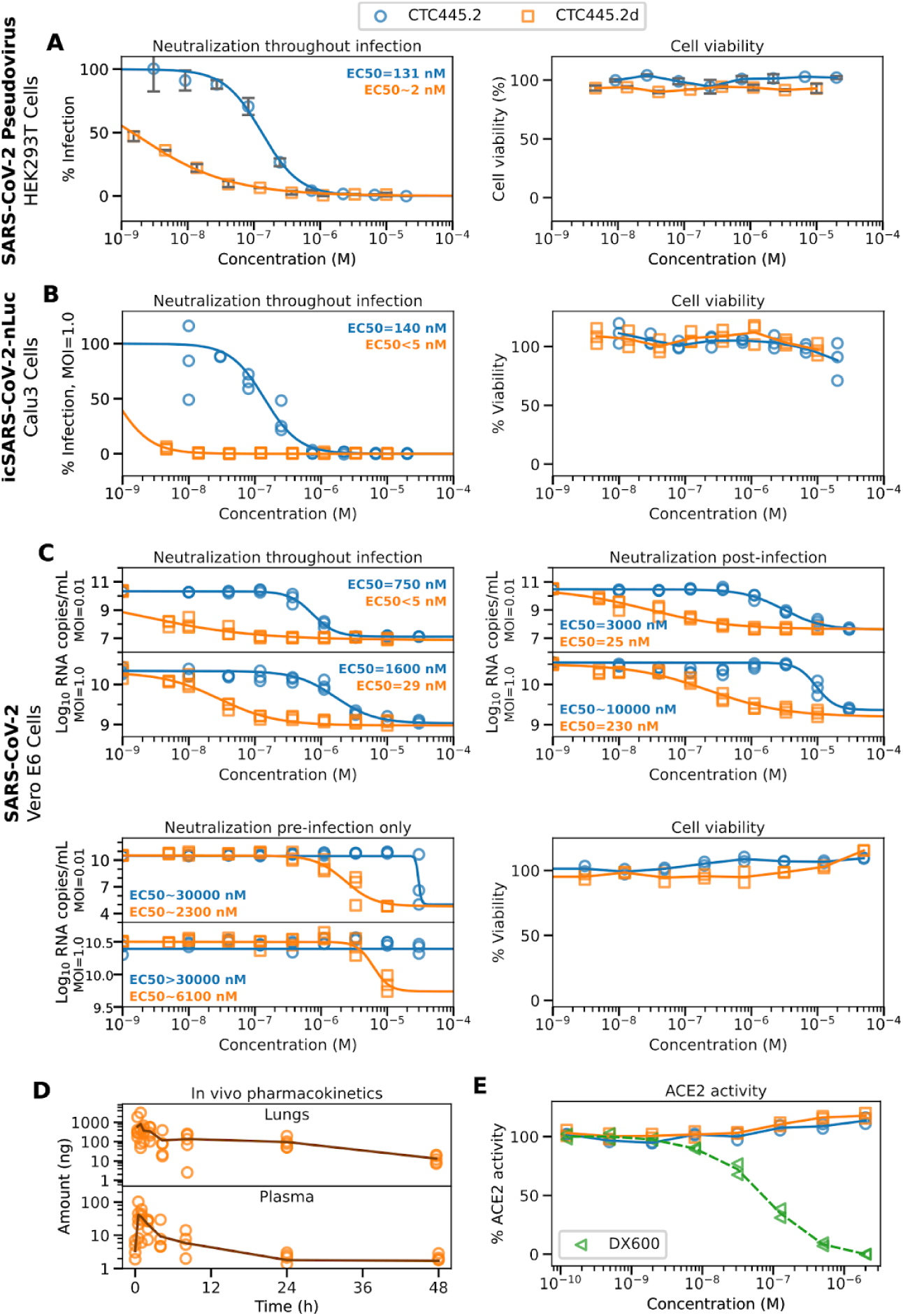
In vitro virus neutralization by CTC-445.2 and CTC-445.2d. CTC-445.2 (blue) and CTC-445.2d (orange) are shown. **A)** SARS-CoV-2/VSV-Luc pseudovirus assay. Neutralization assays (left) were performed using a non-replicative VSV pseudovirus carrying a luciferase reporter gene and expressing the spike protein of SARS-CoV-2 on its surface. Viral neutralization was performed on HEK 293T cells overexpressing ACE2. The de novo decoys were pre-incubated with pseudovirus prior to incubation with cells. Samples were tested in duplicate utilizing 3-fold serial dilutions started at 20 μM (CTC-445.2) or 10 μM (CTC-445.2d). A cell viability assay (right) confirmed that the decoys are not cytotoxic to HEK 293T. We used an analogous pseudovirus expressing an unrelated RBD to confirm that their neutralizing activity of the de novo decoys is specific (figure S11); **B)** (left) Neutralization of NanoLuc SARS-CoV-2 by CTC-445 and CTC-445.2 in Calu-3 cells after a 72 h incubation; (right) A cell viability assay (48 h) confirmed that the decoys are not cytotoxic to Calu-3; **C)** Neutralization of live BetaCoV/Hong Kong/VM20001061/2020 SARS-CoV-2 virus in Vero E6 cells. The cells were incubated with virus at both MOI 1.0 and MOI 0.01 throughout infection (pre-incubated with virus, during infection, and post infection) (upper left), post-infection (upper right) and pre-infection only (lower left). Cell viability in Vero E6 cells (lower right) was independently performed (CCK8 assay) and confirmed that the de novo decoys are not cytotoxic. SARS-CoV-2 RNA copy numbers were determined by quantitative real-time RT-PCR. All assays were performed in triplicate unless otherwise noted; **D)** Bioavailability of CTC-445.2d in mice lung (top) and plasma (bottom) after intranasal administration (see methods); **E)** ACE2 functional activity as measured by enzymatic release of a free fluorophore from Mca-APK(DNP) substrate. The de novo decoys didn’t affect ACE2 function. DX600 was used as positive control (see methods).

### Biophysical and biochemical properties of the *de novo* protein ACE2 decoys

CTC-445 is a 160 a.a. protein comprising 18 of the natural amino acids (it does not contain cysteine or tryptophan residues), has nanomolar affinity for the RBD of SARS-CoV-2 (K_d_ = 357 nM, see table S2 and methods) and can outcompete binding of the SARS-CoV-2 RBD to hACE2 (IC_50_ = 1675 nM, figure 1I). Notably, CTC-445 exhibits lower binding potency for SARS-CoV-2 than hACE2 (K_d_ = 4.6 nM, IC_50_ = 10.9 nM, table S3) and a very weak cross reactivity to SARS-CoV-1 (K_d_ ≈55 µM, figure 2F, table S2 and methods). We believe that the low binding affinity observed is likely due to the low stability of its folded state (ΔG_NI_∼ -2.7 kcal mol-1, T_m_∼ 75.3 °C, figure 1F and 2C-E). In contrast, the experimentally stability-optimized de novo decoy CTC-445.2 is predominantly monomeric (figure 1H and S7), thermodynamically hyperstable (ΔG_NI_∼ -5.0 kcal mol-1, T_m_ ∼ 93°C, figure 2C-D), exhibits low nanomolar affinity for the RBD of SARS-CoV-2 (K_d_ ∼ 21.0 nM, see table S3), and has improved cross-reactivity to SARS-CoV-1 (K_d_ ∼ 7.1 µM, table S2 and figure 2F). CTC-445.2 is capable of outcompeting hACE2 binding to the SARS-CoV-2 RBD at low nanomolar concentrations (IC_50_ ∼ 10.4 nM, figure 1I). Single-site saturation mutagenesis (SSM) (*28, 29*) of the SARS-CoV-2 RBD binding interface showed that the effects of mutations in the binding of hACE2 are closely recapitulated by CTC-445.2 (see figure S14).The bivalent de novo protein decoy CTC-445.2d is our top optimized variant due to its remarkable binding and neutralizing potency, albeit it has a slightly less stable native state than its parent (ΔG_NI_ ∼ -4.5, T_m_ ∼ 71.7°C; figure 2C-D) highlighting perhaps the advantage of starting with a very stable molecule before attempting a domain duplication to augment binding potency. The increased avidity of CTC-445.2d confers a ∼10-fold improvement in binding affinity of the de novo decoy for both SARS-CoV-2 RBD (K_d_ ∼ 3.5 nM, table S2) and SARS-CoV-1 RBD (K_d_ ∼ 587 nM, table S2 and figure S9), as well as a similar increase in its ability to outcompete hACE2 binding to the SARS-CoV-2 RBD (IC_50_ ∼ 0.7 nM, figure 1I).

### Functional characterization of CTC-445.2 and CTC-445.2d

The strong binding of our optimized *de novo* protein decoys to the SARS-CoV-2 RBD translated into effective and specific*in vitro*neutralization of viral cell infection (figure 3). The presence of the *de novo* decoys showed no impact on mammalian cell viability. Co-incubation of hACE2 with the decoys had no impact on the enzymatic activity of hACE2 (figure 3E). Both of the *de novo* hACE2 decoys were able to fully neutralize viral infection in three different *in vitro* systems (figure 3A-C). Briefly, in a VSV pseudovirus system expressing the SARS-CoV-2 spike protein, the *de novo* decoys specifically protected HEK 293T cells overexpressing hACE2 from infection (figure 3A and figure S11). Our *de novo* protein decoys also prevented infection by SARS-CoV-2 expressing the nanoLuc reporter (SARS-CoV-2 nanoLuc, see methods) in the lung epithelial cell line Calu-3, which expresses both ACE2 and the transmembrane protease serine 2 (TMPRSS2), known to augment the infectivity of SARS-CoV-2 (*30, 31*) (figure 3B; EC_50_ < 5 nM at MOI=1.0). To confirm the mechanism of inhibition (i.e. extracellular neutralization of the virus) by CTC-445.2 and CTC-445.2d, a time-of-addition assay was performed in Vero E6 cells. Briefly, CTC-445.2 or CTC-445.2d were added either during the full course of infection (i.e. co-incubated with the virus and in the cell media), before infection (i.e. only co-incubated with the virus), or after infection (i.e. only in the cell media). As expected, CTC-445.2 and CTC-445.2d were most effective when continuously present throughout the full course of infection (figure 3C and S12). To define the activity of CTC-445.2d as a locally delivered therapeutic to possibly protect the airway from infection, we delivered a high dose (100 µg) of CTC-445.2d to Balb/c mice via inhalation of intranasal droplets. We observed persistence of fully functional CTC-445.2d in the lungs for more than 24 hours (figure 3D). We also detected CTC-445.2d in blood, raising the possibility that inhaled therapy might lead to some level of systemic exposure.

## Conclusions

The emergence of SARS-CoV-2 as a global health threat has highlighted the need for the rapid development of novel countermeasures. Soluble “decoy” proteins are orthogonal to traditional therapeutic approaches such as vaccination, neutralizing antibodies, or small molecule inhibitors, and might have the potential to better overcome the problem of mutational viral evasion. Utilizing natural proteins as therapeutics often presents a number of significant challenges, such as poor stability that can complicate manufacturing, transport and storage, side effects that can arise from residual/unwanted biological activity, and the fact that human proteins repurposed as therapeutics pose the risk of eliciting an autoimmune response with possibly lifelong consequences (*32–41*). Designed *de novo* protein decoys offer an avenue to overcome these challenges. Because CTC-445.2d differs in sequence and structure from any natural protein, it reduces the potential risk of eliciting an autoimmune reaction. The optimized *de novo* protein decoys are easy to manufacture in traditional bacterial systems, and their thermodynamic hyperstability and full thermal reversibility make them ideal candidates for transport, storage, and local delivery. Also, due to their remarkable thermodynamic stability, there is further potential to fine tune *de novo* protein decoys (*42*). For example, we have shown thatby simply increasing the avidity of CTC-445.2 it is possible to greatly augment its potency. As well, we demonstrate that it is also possible to further increase stability while preserving the function of the de novo decoy (figure S13). This tunability could be difficult to achieve with a natural protein such as hACE2, which is much larger (∼90 kDa) and has significantly lower stability (figure S8).The protein surface that CTC-445 (and its optimized variants) present toSARS-CoV-2 is designed to be virtually identical to the natural binding surface that the virus targets in hACE2. Therefore, mutations in the RBD that affect binding to our de novo decoy are likely to have a similar effect on the virus’ binding to hACE2, effectively making the decoys functionally resilient to mutational escape (figure S10 and S14). Our principle is also exemplified by the cross reactivity of our de novo decoys with SARS-CoV-1 despite (at least) 17 mutations that separate their binding interfaces (figure S9), including a drastic loop remodeling in the RBD’s ridge region (*43*).

We have laid the groundwork for a generalized strategy that has the potential to quickly address emerging biological threats. Our strategy is novel, precise, and fast. CTC-445.2d is, to our knowledge, the first reported example of a *de novo* protein therapeutic that has been conceptualized, engineered, and validated in less than ten weeks, matching the rapid speed with which the SARS-CoV-2 threat has emerged. Moreover, further speed and accuracy improvements to our strategy are theoretically possible (figure 1G). We envision that future embodiments of our *de novo* protein design paradigm will be generally useful for the rapid development of therapeutic decoys to treat disease, not only for viral agents, but for any case where interrupting protein-protein interactions could be beneficial. Combined prowess in structural biology and *de novo* protein design are quickly bringing us to the point where therapeutic molecules can be engineered on demand.

## Acknowledgments

The authors thank Michael Dougan, Leslie Aberman, Umut Ulge, Julie Rathbun, and Jonathan Drachman, for useful discussions and comments on this manuscript; Neoleukin Therapeutics, Inc. (“Neoleukin”) for supporting this work. All the computational resources for the *de novo* protein design were provided by Neoleukin’s high-performance “Neo” computational cluster. NIH grants AI145296 and AI127463 to M.G.Jr.; HHSN272201400006C from NIAID, NIH to H.-L.Y and R01AI108197 to RSB. Neoleukin™ is a trademark of Neoleukin. The views and opinions expressed in this article are those of the authors and do not necessarily reflect the position of Neoleukin.

## Author Contributions

T.W.L. designed and coordinated the research, developed computational design methods, designed *de novo* protein decoys of ACE2, characterized designs, and wrote the manuscript; R.V. designed *de novo* proteins, performed molecular biology, characterized and optimized the designs, and wrote the manuscript; N.C. designed *de novo* proteins, characterized and optimized the designs, and wrote the manuscript; J.W.N. designed *de novo* proteins, characterized and optimized the designs, performed molecular biology, performed SSM experiments, and wrote the manuscript; M.J.W. designed *de novo* proteins, performed molecular biology, characterized and optimized the designs, and wrote the manuscript; W.S. performed cell-neutralization assays with the live SARS-CoV-2 virus in Vero E6 cells and edited the manuscript; T.-Y.H. performed cell-neutralization assays with the live SARS-CoV-2 NanoLuc virus in Calu-3 cells; K.E.-N. performed cell-neutralization assays with the live SARS-CoV-2 NanoLuc virus in Calu-3 cells; YJH developed the nLUC reporter virus; K.Y. designed and performed ACE2 competition assays and developed methods to quantify the *de novo* designs in tissue lysates; T.P. designed *de novo* proteins and purified and characterized the *de novo* proteins; M.M. designed *de novo* proteins; A.P. designed *de novo* proteins and performed binding characterizations; U.Y.L. designed *de novo* proteins; M.Ma. performed pharmacokinetic studies in mice; J.C. performed pharmacokinetic studies in mice; A.C. performed ACE2 enzymatic assay and cytotoxicity assays with VeroE6; T.B. purified and characterized the *de novo* proteins; H.P. performed mass spectrometry; N.S.C. performed molecular biology; J.Ca. developed and implemented computational tools for collaborative *de novo* protein design; Y.-R.L. designed *de novo* proteins; A.J.-D. coordinated project operations and wrote the manuscript; RSB coordinated the development of the nLUC reporter virus and edited the paper; C.D.W. coordinated the research for ACE2 competition assays and methods to quantify the *de novo* designs and edited the paper; R.S. coordinated the research for in vitro neutralization testing and in vivo pharmacokinetics of the *de novo* proteins and edited the paper; M.G.Jr. coordinated and directed the research for in vitro NanoLuc SARS-CoV-2 neutralization and edited the paper; L.M.B.-M. designed *de novo* proteins, coordinated the purification and characterization of the *de novo* proteins and edited the paper; H.-L.Y. coordinated the research for in vitro SARS-CoV-2 neutralization and edited the paper; D.-A.S. generated the original idea to design the *de novo* decoys to neutralize SARS-CoV-2, designed the research, developed computational design selection strategies, wrote the manuscript, and directed the effort.

## List of supplementary materials

1. **Materials and methods**
2. **Table S1-S3** **Table S1**. SARS-CoV-2 neutralization by CTC-445.2 and CTC-445.2d in a time-of-addition experiment. **Table S2**. Characterization of recombinant, purified CTC-445 variants. **Table S3**. Binding kinetics of de novo designed decoys with S protein RBDs.
3. **Supplementary figures S1-S12:** **Figure S1**. Target mechanism of inhibition of SARS-CoV-2 by the de novo designed ACE2 decoys. **Figure S2**. Three dimensional structures of ACE2 in-complex with SARS-CoV-1 and SARS-CoV-2 RBD. **Figure S3**. Yeast display characterization of CTC-445 binding to SARS-CoV-2 RBD and SARS-CoV-1 RBD. **Figure S4**. Sequence alignment for the directed evolution of CTC-445 using error-prone PCR and yeast display FACS. **Figure S5**. Potency of CTC-445 variants *v*.*s*. its molecular weight. **Figure S6**. Mass spectrometry of CTC-445.2 and CTC-445.2d. **Figure S7**. SEC-MALS of de novo designed decoys CTC-445, CTC-445.2 and CTC-445.2d. **Figure S8**. Thermal stability of hACE2. **Figure S9**. Binding kinetics of CTC-445.2 and CTC-445.2d to SARS-CoV-1 RBD. **Figure S10**. Kinetics of binding for CTC-445.2 to SARS-CoV-2 RBD mutants. **Figure S11**. Inhibition of infection by VSV-Luc pseudovirus expressing VSVg. **Figure S12**. Inhibition of SARS-CoV-2 (cytopathic effect) in Vero E6 cells. **Figure S13**. CTC-445.3d, a further optimized CTC-445.2d variant. Stability, thermal kinetics, binding and in vitro neutralization. **Figure S14**. Yeast-display single-site saturation mutagenesis (SSM) library of the SARS-CoV-2 RBD binding interface, effect on binding of CTC-445.2 and hACE2.

