## Supplemental Material for "*De novo* design of ACE2 protein decoys to neutralize SARS-CoV-2"

**List of supplementary materials**

1. **Materials and methods**
2. **Table S1-S3**

**Table S1.** SARS-CoV-2 neutralization by CTC-445.2 and CTC-445.2d in a time-of-addition experiment.

**Table S2.** Characterization of recombinant, purified CTC-445 variants.

**Table S3.** Binding kinetics of de novo designed decoys with S protein RBDs.

3. **Supplementary figures S1-S12:**

**Figure S1**. Target mechanism of inhibition of SARS-CoV-2 by the de novo designed ACE2 decoys.

**Figure S2.** Three dimensional structures of ACE2 in-complex with SARS-CoV-1 and SARS-CoV-2 RBD.

**Figure S3.** Yeast display characterization of CTC-445 binding to SARS-CoV-2 RBD and SARS-CoV-1 RBD.

**Figure S4.** Sequence alignment for the directed evolution of CTC-445 using error-prone PCR and yeast display FACS.

**Figure S5.** Potency of CTC-445 variants *v.s.* its molecular weight.

**Figure S6.** Mass spectrometry of CTC-445.2 and CTC-445.2d.

**Figure S7.** SEC-MALS of de novo designed decoys CTC-445, CTC-445.2 and CTC-445.2d.

**Figure S8.** Thermal stability of hACE2.

**Figure S9.** Binding kinetics of CTC-445.2 and CTC-445.2d to SARS-CoV-1 RBD.

**Figure S10.** Kinetics of binding for CTC-445.2 to SARS-CoV-2 RBD mutants.

**Figure S11.** Inhibition of infection by VSV-Luc pseudovirus expressing VSVg.

**Figure S12.** Inhibition of SARS-CoV-2 (cytopathic effect) in Vero E6 cells.

**Figure S13.** CTC-445.3d, a further optimized CTC-445.2d variant. Stability, thermal kinetics, binding and in vitro neutralization.

**Figure S14.** Yeast-display single-site saturation mutagenesis (SSM) library of the SARS-CoV-2 RBD binding interface, effect on binding of CTC-445.2 and hACE2.

### Supplementary Materials

**Table S1: SARS-CoV-2 neutralization by CTC-445.2 and CTC-445.2d in a time-of-addition experiment.** *Recombinant protein was present before (pre-incubating with virus for 1h), during (1h), and post SARS-CoV-2 infection (72h) in Vero E6 cells. #Recombinant protein was added post SARS-CoV-2 infection. @Recombinant protein was pre-incubated with virus for 1h and the virus-recombinant protein mix was added to Vero E6 cells and incubated for 1h. The cells were washed with PBS after infection and overlaid with media without recombinant protein.

| Proteins | Inhibitory concentration at MOI= 1.0 (µM) | | | Inhibitory concentration at MOI= 0.01 (µM) | |
|---|---|---|---|---|---|
| | CPE formation | Reduction in viral RNA copy numbers ($EC_{50}$) | | CPE formation | Reduction in viral RNA copy numbers ($EC_{50}$) |
| Experiment 1: Inhibitory effect throughout the course of infection* | | | | | |
| CTC-445.2 | 10 | 1.807 | | 2.31 | 0.753 |
| CTC-445.2d | 0.37 | 0.0235 | | 0.01 | <0.005 |
| Experiment 2: Post-infection inhibitory effect# | | | | | |
| CTC-445.2 | 30 | ~10 | | 10 | 2.947 |
| CTC-445.2d | 3.33 | 0.2369 | | 0.12 | 0.0139 |
| Experiment 3: Pre-infection inhibitory effect@ | | | | | |
| CTC-445.2 | >30 | >30 | | ~30 | ~29.33 |
| CTC-445.2d | >10 | ~3.903 | | 3.33 | 2.688 |

**Table S2: Characterization of recombinant, purified CTC-445 variants.** CTC-445 variants were generated by rationally combining mutations discovered by directed evolution and characterized. Columns indicate the qualitative expression level (0-5), solubility (0-5), and binding affinity (0-5). Propensity for aggregation indicates whether large molecular weight aggregates or oligomers were present during size exclusion chromatography, and the fraction of non-monomeric protein is measured by comparing peak areas for monomeric and oligomeric species by analytical size exclusion chromatography.

| ID | Description | Expression level (Cat. 0-3) | Solubility (Cat. 0-3) | Binding strength (Cat. 0-5) | Propensity for aggregation | SEC monomeric fraction (%) |
|---|---|---|---|---|---|---|
| CTC-00445 | CTC-445 | | | | | |
| CTC-00613 | CTC-445_S1P | 2 | 1 | 1 | | |
| CTC-00614 | CTC-445_M6L | 3 | 2 | 3 | | |
| CTC-00615 | CTC-445_F152W | 3 | 2 | 1 | | |
| CTC-00616 | CTC-445_F11Y | 3 | 2 | 1 | | |
| CTC-00617 | CTC-445_F11K | 2 | 3 | 2 | | |
| CTC-00618 | CTC-445_R42S | 2 | 1 | 1 | | |
| CTC-00619 | CTC-445_E46S | 2 | 2 | 1 | | |
| CTC-00620 | CTC-445_A88N | 3 | 2 | N.D. | | |
| CTC-00621 | CTC-445_F116N | 2 | 3 | 1 | | |
| CTC-00622 | CTC-445_R124S | 3 | 3 | 2 | | |
| CTC-00623 | CTC-445_F137V | 3 | 3 | N.D. | | |
| CTC-00624 | CTC-445_R143L | 2 | 3 | 3 | | |
| CTC-00625 | CTC-445_S1P_F11Y_R42S_E46S_A88N_F116N_R124S_F137V_R143L | 3 | 3 | N.D. | | |
| CTC-00626 | CTC-445_S1P_M6L_F11Y_R42S_E46S_A88N_F116N_R124S_F137V_R143L_F152W | 2 | 3 | 0.5 | | |
| CTC-00627 | CTC-445_S1P_F11K_R42S_E46S_A88N_F116N_R124S_F137V_R143L | 2 | 3 | N.D. | | |
| CTC-00628 | CTC-445_S1P_M6L_F11K_R42S_E46S_A88N_F116N_R124S_F137V_R143L_F152W | 2 | 3 | 2 | | |
| CTC-00629 | CTC-445_E74C | 3 | 2 | 1 | | |
| CTC-00630 | CTC-445_K63C | 2 | 1 | Not screened | | |
| CTC-00631 | CTC-445_F11C | 2 | 1 | N.D. | | |
| CTC-00632 | CTC-445_E74C_F11Y | 2 | 3 | 1 | 0 | 17% |
| CTC-00633 | CTC-445_K63C_F11Y | 2 | 1 | 1 | 1 | |

| | | | | | | |
|---|---|---|---|---|---|---|
| CTC-00634 | CTC-445_E74C_S1P_M6L_F11Y_R42S_E46S_A88N_F116N_R124S_F137V_R143L_F152W | 2 | 3 | N.D. | 0 | 82% |
| CTC-00635 | CTC-445_K63C_S1P_M6L_F11Y_R42S_E46S_A88N_F116N_R124S_F137V_R143L_F152W | 2 | 3 | N.D. | 0 | 86% |
| CTC-00636 | CTC-445_S1P_M6L_F11C_R42S_E46S_A88N_F116N_R124S_F137V_R143L_F152W | 0 | 1 | N.D. | 1 | |
| CTC-00637 | {PG}.CTC-445_M6L_R126L | 1.5 | 2 | 4 | 0 | 97% |
| CTC-00638 | {PG}.CTC-445_F116L_R124S | 0.5 | 3 | Not screened | 1 | |
| CTC-00639 | {PG}.CTC-445_M6L_F116L_R124S_R126L | 2 | 2 | 4.5 | 0 | 92% |
| CTC-00640 (CTC-445.2) | {PG}.CTC-445_M6L_E86G_F116L_R124S_R126L | 2 | 3 | 4.5 | 0 | 99% |
| CTC-00641 | {PG}.CTC-445_A2V_E3V_M6L_I61V_K63R_A73V_K84T_E86G_A92V_A95E_K100Q_F116L_R124S_R126L_A136T_R143S | 2 | 3 | 3.5 | 0 | 95% |
| CTC-00642 | {PG}.CTC-445_M6L_F11C_R126L | 3 | 2 | 4.5 | 1 | |
| CTC-00643 | {PG}.CTC-445_M6L_K63C_R126L | 3 | 2 | 4.5 | 0 | 59% |
| CTC-00644 | {PG}.CTC-445_M6L_E74C_R126L | 3 | 2 | 4.5 | 0 | 89% |
| CTC-00645 | {PG}.CTC-445_F11C_F116L_R124S | 3 | 2 | Not screened | | |
| CTC-00646 | {PG}.CTC-445_K63C_F116L_R124S | 3 | 2 | 3.5 | 1 | |
| CTC-00647 | {PG}.CTC-445_E74C_F116L_R124S | 3 | 2 | 3 | 1 | |
| CTC-00648 | {PG}.CTC-445_M6L_F11C_F116L_R124S_R126L | 2 | 3 | 5 | 0 | 85% |
| CTC-00649 | {PG}.CTC-445_M6L_K63C_F116L_R124S_R126L | 2 | 3 | 4.5 | 0 | 87% |
| CTC-00650 | {PG}.CTC-445_M6L_E74C_F116L_R124S_R126L | 0.5 | 3 | 4.5 | 0 | 90% |
| CTC-00651 | {PG}.CTC-445_A2V_E3V_M6L_F11C_I61V_K63R_A73V_K84T_E86G_A92V_A95E_K100Q_F116L_R124S_R126L_A136T_R143S | 2 | 3 | 4 | 0 | 97% |

| | | | | | | |
|---|---|---|---|---|---|---|
| CTC-00652 | {PG}.CTC-445_A2V_E3V_M6L_I61V_K63R_K63C_A73V_K84T_E86G_A92V_A95E_K100Q_F116L_R124S_R126L_A136T_R143S | 2 | 3 | 4 | 0 | 89% |
| CTC-00653 | {PG}.CTC-445_A2V_E3V_M6L_I61V_K63R_A73V_E74C_K84T_E86G_A92V_A95E_K100Q_F116L_R124S_R126L_A136T_R143S | 2 | 3 | 4 | 0 | 86% |
| CTC-00654 **(CTC-445.2d)** | {PG}.CTC-445_M6L_E86G_F116L_R124S_R126L_X2 | 2 | 3 | 5 | | |
| CTC-00655 | {PG}.CTC-445_A2V_E3V_M6L_I61V_K63R_A73V_K84T_E86G_A92V_A95E_K100Q_F116L_R124S_R126L_A136T_R143S_X2 | 2 | 3 | 5 | | |

**Table S3. Binding kinetics of de novo designed decoys with S protein RBDs.**

| Decoy | RBD | $k_{on}$ ($M^{-1}$ $s^{-1}$) | $k_{off}$ ($s^{-1}$) | $K_d$ kinetics (nM) | $K_d$ steady-state (nM) |
|---|---|---|---|---|---|
| hACE2 | SARS-Cov-1 | $6.8 \times 10^4$ | $1.0 \times 10^{-3}$ | 14 | 43 |
| | SARS-CoV-2 | $8.6 \times 10^4$ | $4 \times 10^{-4}$ | 4.7 | 31 |
| CTC-445 | SARS-CoV-1 | ~ $4.5 \times 10^4$ | ~$2.5 \times 10^{-1}$ | ~ $5.5 \times 10^4$ | ND |
| | SARS-CoV-2 | $8.6 \times 10^4$ | $5.5 \times 10^{-2}$ | 646 | 357 |
| CTC-445.2 | SARS-CoV-1 | $1.0 \times 10^5$ | $3.9 \times 10^{-1}$ | $3.8 \times 10^3$ | $7.1 \times 10^3$ |
| | SARS-CoV-2 | $3.9 \times 10^5$ | $1.2 \times 10^{-2}$ | 30 | 21 |
| | SARS-CoV-2_N354D | $5.0 \times 10^5$ | $1.5 \times 10^{-2}$ | 30 | 22 |
| | SARS-CoV-2_N354D_D364Y | $7.4 \times 10^5$ | $1.5 \times 10^{-2}$ | 20 | 16 |
| | SARS-CoV-2_V367F | $5.9 \times 10^5$ | $1.3 \times 10^{-2}$ | 22 | 17 |
| | SARS-CoV-2_R408I | $5.1 \times 10^5$ | $1.7 \times 10^{-2}$ | 33 | 26 |
| | SARS-CoV-2_W436R | $6.8 \times 10^5$ | $8.7 \times 10^{-2}$ | 13 | 8 |
| CTC-445.2d | SARS-CoV-1 | $3.7 \times 10^5$ | $4.7 \times 10^{-1}$ | $1.2 \times 10^3$ | 587 |
| | SARS-CoV-2 | $3.0 \times 10^5$ | $\leq 2.0 \times 10^{-3}$ | $\leq 7.0$ | 3.5 |

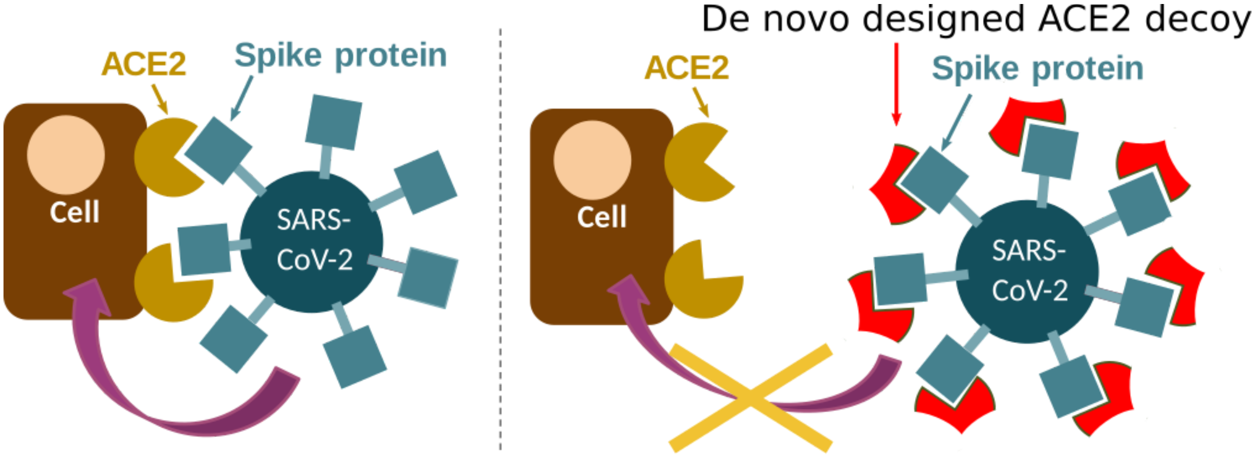


**Figure S1. Target mechanism of inhibition of SARS-CoV-2 by the de novo designed ACE2 decoys.** SARS-CoV-2 (and SARS-CoV) gains entry into cells by first binding to the ACE2 receptor on the surface of human cells via the spike protein (left). The spike protein binds to the de novo designed ACE2 decoys using the same binding interface as natural ACE2, effectively making decoys functionally resilient to mutation. The SARS-CoV-2 spike protein binds to the decoys instead of native ACE2, sequestering the virus and preventing viral entry into cells while keeping native ACE2 function intact (right).

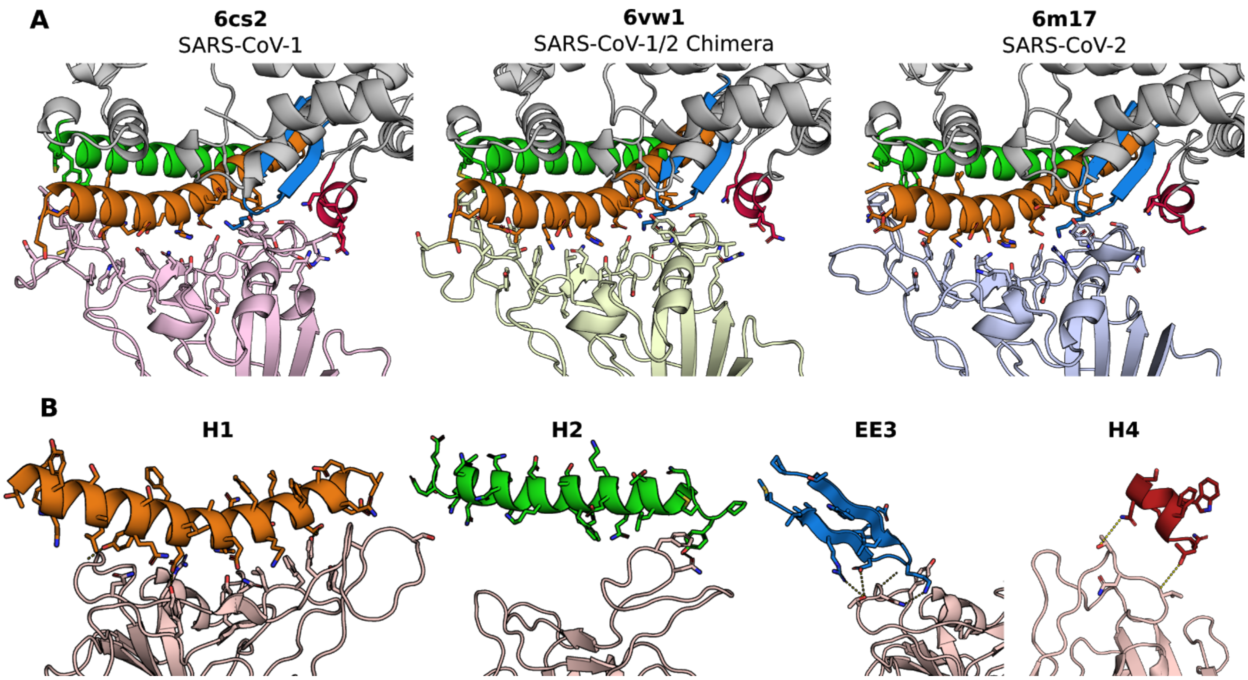


**Figure S2. Three dimensional structures of ACE2 in-complex with SARS-CoV-1 and SARS-CoV-2 RBD. A)** Comparison of ACE2 models used to identify motifs for designed proteins. H1 (orange), H2 (green), EE3 (blue), and H4 (red) are the motifs that interact with the RBD of SARS-CoV-1 (pink, left), a SARS-CoV-1+SARS-CoV-2 chimera (light green, middle) and SARS-CoV-2 (light blue, right). Computational designs were created against one of these structures, but evaluated against the S protein RBD from all three RBDs. **B)** Structural motifs involved in binding of spike protein to ACE2 and their interactions with SARS-CoV-2 spike protein. H1 and H2 were present in all designs, and EE3 was present in the majority of designs. To simplify the design process of the de novo decoys, we did not incorporate the fourth binding motif (H4) in our design strategy because we predicted it to have a minor contribution to binding.

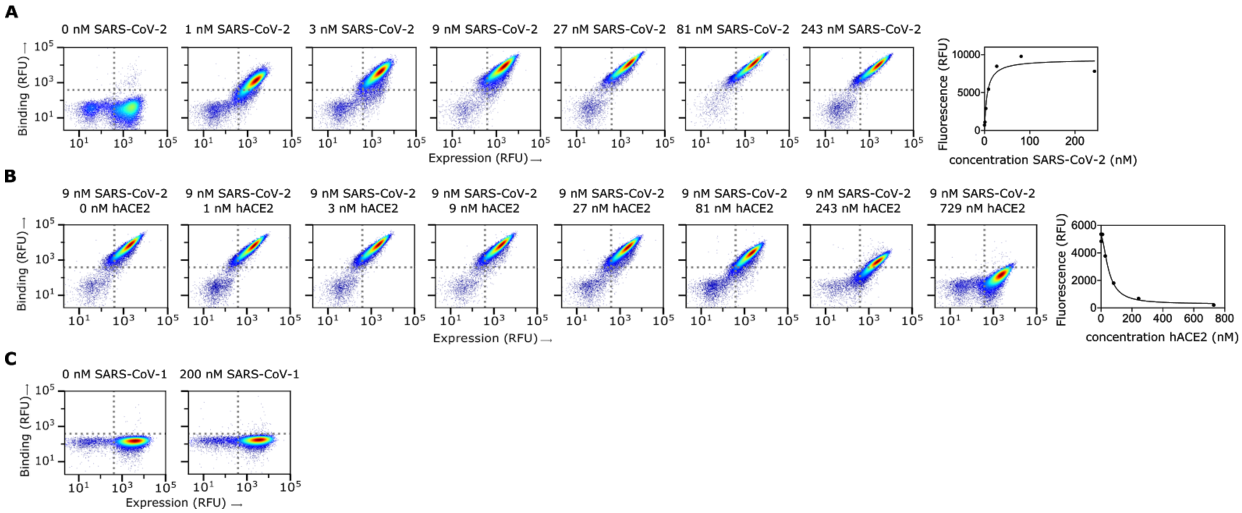


**Figure S3. Yeast display characterization of CTC-445 binding to SARS-CoV-2 RBD and SARS-CoV-1 RBD. A)** Titration assay to estimate the binding $K_d$ of CTC-445 for SARS-CoV-2 RBD. Yeast displaying CTC-445 were incubated with increasing concentrations of SARS-CoV-2 RBD (0, 1, 3, 9, 27, 81 and 243 nM, see methods). On the right, steady state plot to reveal a $K_d$ of 6 nM. **B)** Competition assay to estimate the $IC_{50}$ of hACE2 to compete binding of CTC-445 to SARS-CoV-2 RBD. Yeast displaying CTC-445 were incubated with SARS-CoV-2 9 nM and increasing concentrations of hACE2 (0, 1, 3, 9, 27, 81, 243 and 729 nM, see methods). On the right, steady state plot to reveal an $IC_{50}$ of 49 nM. **C)** Screen for binding of CTC-445 to SARS-CoV-1 RBD. Yeast displaying CTC-445 were incubated with SARS-CoV-1 RBD at 200 nM (see methods), to reveal no binding at this concentration.

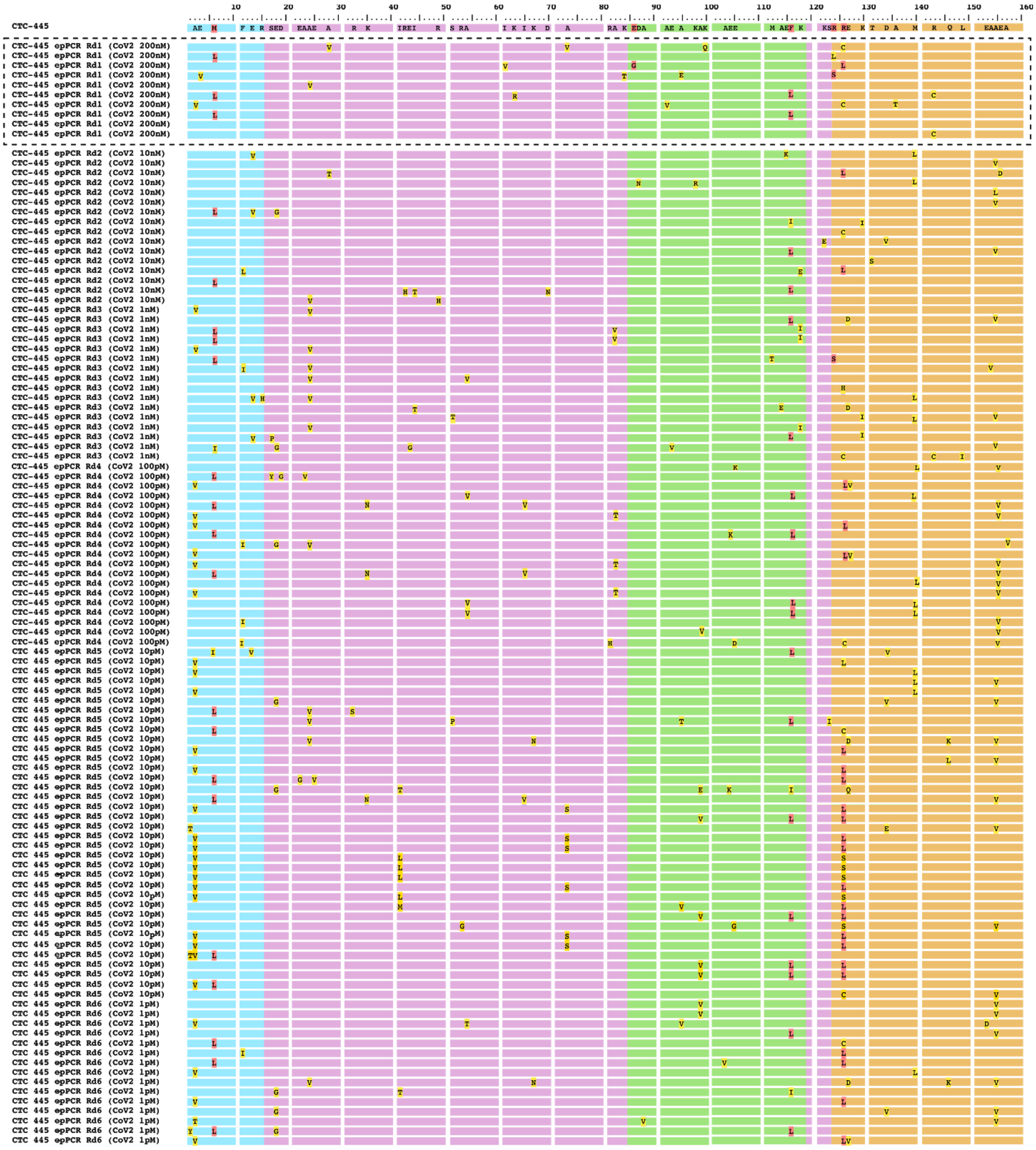

**Figure S4. Sequence alignment for the directed evolution of CTC-445 using error-prone PCR and yeast display FACS.** Error Prone PCR was performed on the gene coding for CTC-445 to identify mutations that improve its stability and affinity to SARS-CoV-2. A DNA library of 2x10$^6$ CTC-445 variants was generated and transformed into yeast for yeast display. Six rounds of yeast sorting were performed, with increasingly stringent binding conditions. For every round of cell sorting, the CTC-445 library was first treated with a mixture of trypsin and chymotrypsin (see methods), washed, and then allowed to bind to SARS-CoV-2 RBD at concentrations of: 200 nM for round 1, 10 nM for round 2, 1 nM for round 3, 100 pM for round 4, 10 pM for round 5 and 1 pM for round 6 (see methods). The top 5% binders for every round were sorted using fluorescence-activated cell sorting (FACS) and selected cells were plated on selective SDCAA plates for sequencing. Cartoon representations of the sequences are shown here with H1 in orange, H2 in green, EE3 in blue and supporting motifs in magenta. CTC-445 is shown on the top and positions where mutations were found are indicated with the residue letter. Ten sequences were picked in the first round of sorting, highlighted with a dashed box. Mutations found in round 1 in combination with rational design led to the CTC-445 second generation variants in Table S2, which includes CTC-445.2. CTC-445.2 is five mutations away from CTC-445 (M6L, E86G, F116L, R124S and R126L, highlighted in red). The more stringent rounds of sorting further confirmed mutations M6L, F116L and R126L and revealed new mutations (e.g. A2V and A155V) that are located in the core of the design and were shown to further improve CTC-445 stability (see figure S13).

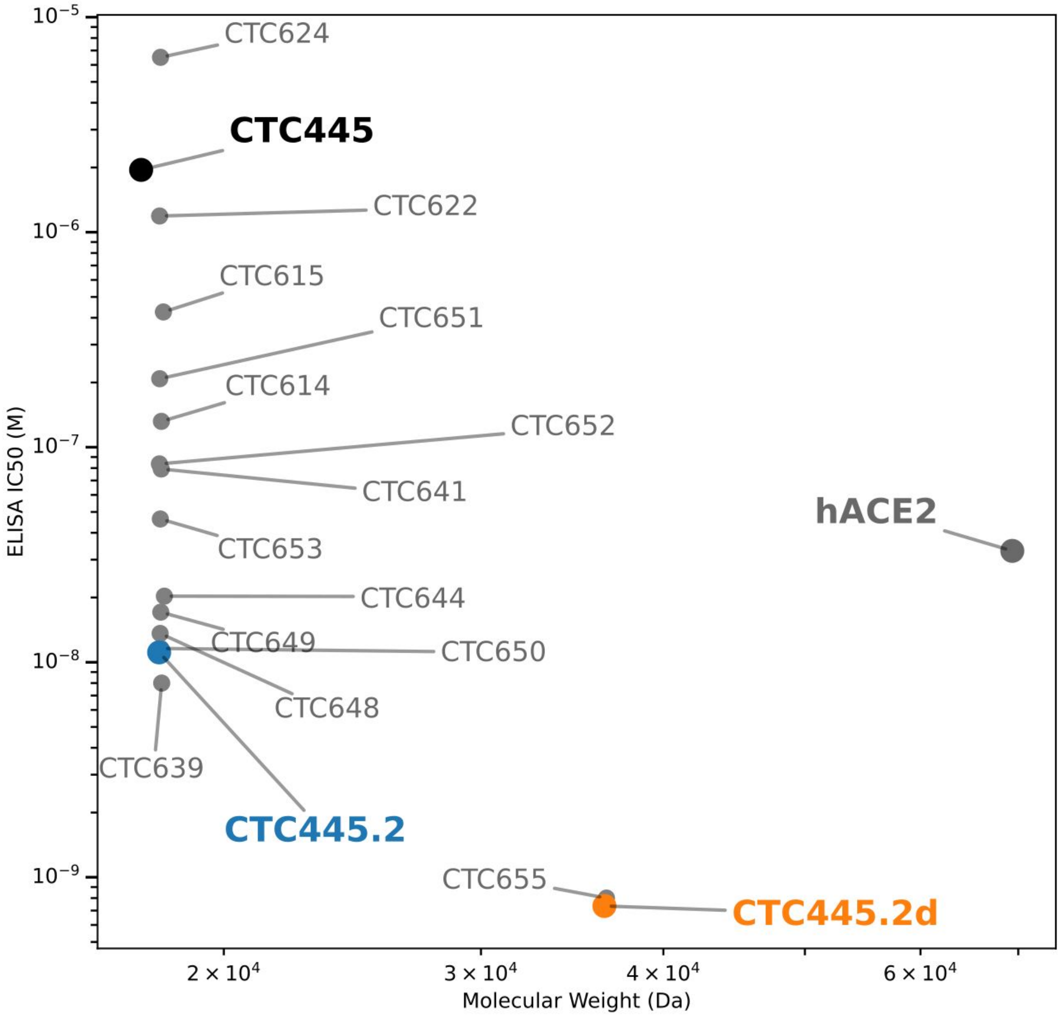


**Figure S5. Potency of CTC-445 variants *v.s.* its molecular weight.** IC50 values for CTC-445 variants for binding to SARS-CoV-2 S/RBD were measured by ELISA in the presence of 0.4 nM biotinylated soluble hACE2. hACE2 bound to RBD was quantified by treatment with streptavidin-HRP and measurement of absorbance at 450 nm.

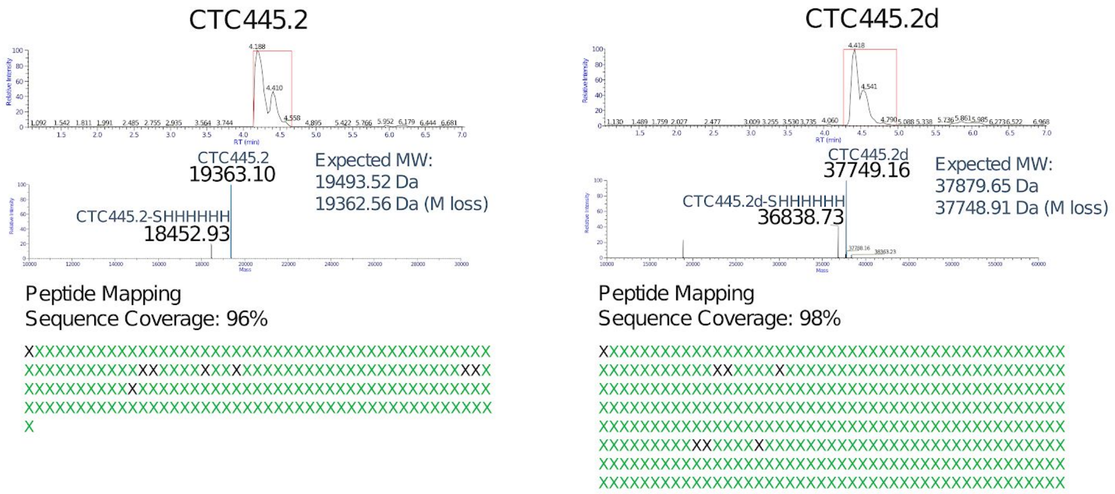


**Figure S6. Mass spectrometry of CTC-445.2 and CTC-445.2d.**

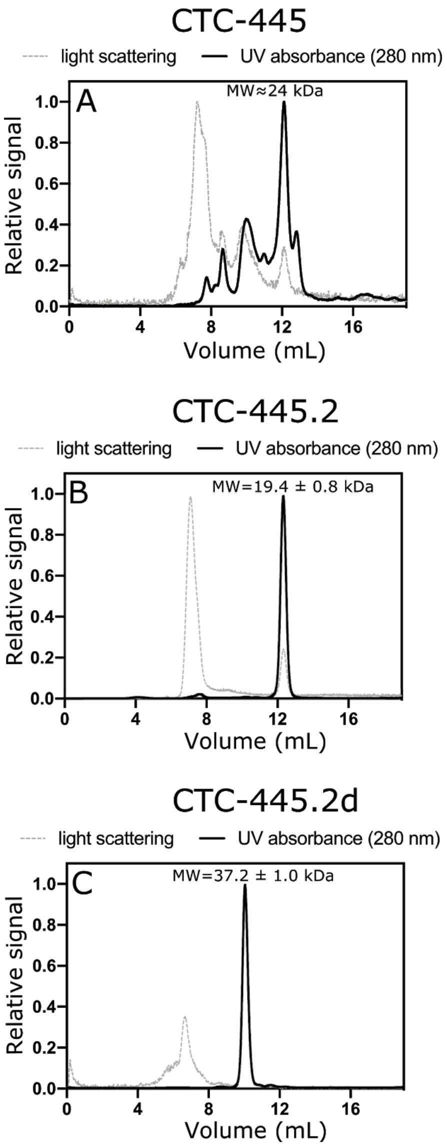


**Figure S7. SEC-MALS analysis of de novo designed decoys CTC-445, CTC-445.2 and CTC-445.2d using a Superdex 75 Increase 10/300 GL size-exclusion column.**
Light scattering signal (gray, dashed) and 280nm UV absorption signal (black, continuous). Chromatogram traces are normalized to the peak with the highest intensity for clarity. Flow rate 0.5 mL/min, injection volume: 100 mL. After isolation of the monomeric CTC-445 **(A)** and CTC-445.2 **(B)** chromatograms shown a small population of high molecular weight aggregates and oligomeric species with high light scattering; while CTC445.2d **(C)** shows evidence of a minuscule population of dimeric oligomer with small light scattering. In all cases chromatograms exhibit excellent monomer-oligomer separation and the light scattering signal exhibits high signal-to-noise ratio. The calculated molecular weights are shown for the fractions of the highest UV signal. For CTC445.2 and CTC445.2d the monomer MW values exhibit high homogeneity and the Mw/Mn ratio was 1.0 which indicates the proteins are monodisperse.

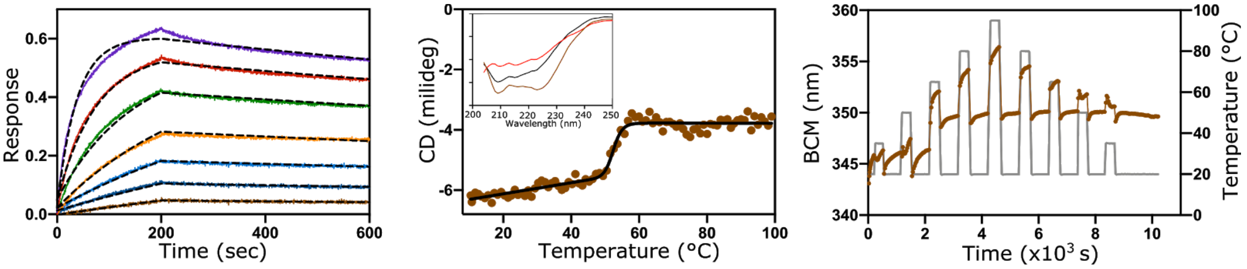


**Figure S8: Binding kinetics and thermal stability of hACE2.** Binding kinetics (left) of hACE2 to immobilized SARS-CoV-2 RBD (300 nM to 4.7 nM); data were globally fit using a 1:1 binding model. Thermal-induced unfolding (middle) measured by circular dichroism at 208 nm was fit using a two-states equation; a $T_m$ value of 52.5 ± 0.5 °C was calculated. The inset shows far UV spectra at 20 °C, 99 °C and 20 °C after heating and cooling the sample. Thermal-recovery assay (right); BCM signal does not return to the initial value after repeated cycles of heating and cooling which indicates poor reversibility.

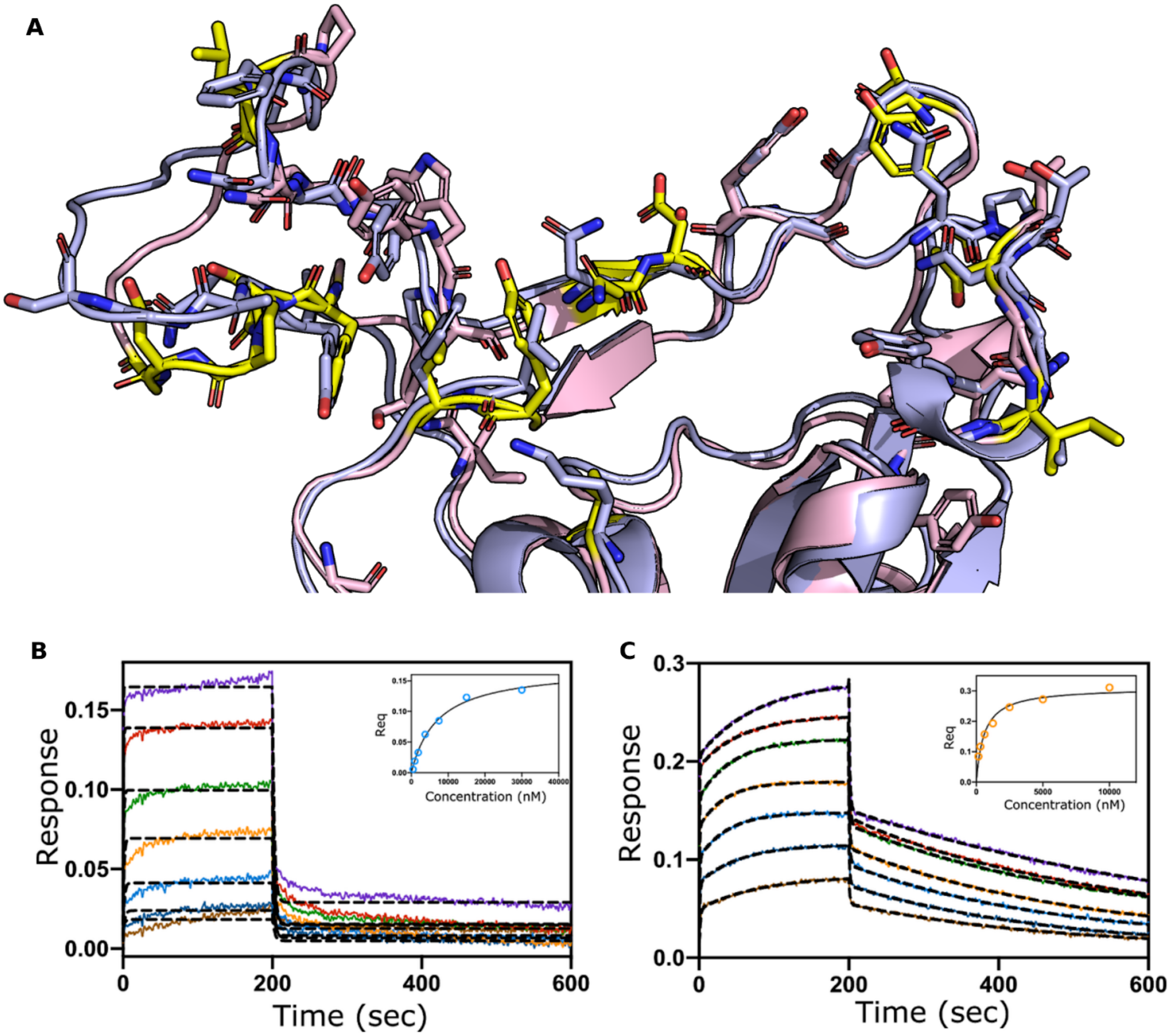


**Figure S9: Binding of CTC-445.2 and CTC-445.2d to SARS-CoV-1 RBD.** A) ACE2-binding interface residues of SARS-CoV (PDB 6CS2; pink) and SARS-CoV-2 (PDB 6M17; blue). The interface (CB-CB distance < 10, sidechains pointing toward surface) contains 29 amino acid positions, 17 of which differ, and a loop that adopts a different conformation. Relative to SARS-CoV-2 spike protein, the interface mutations in the SARS-CoV-1 spike protein (yellow) are: K417V, G446T, L455Y, F456L, Y473F, Q474S, A475P, G476D, S477G, G485A, F486L, Q493N, S494D, Q498Y, P499T, N501T, and V503**I**. B/C) CTC-445.2 (B) and CTC-445.2d (C) were in solution against immobilized SARS-CoV-1 S RBD. The results were globally fit as described above for CTC-445.2 ($K_d$=3.8 μM, $k_{on}$=1.0 x$10^5$ $M^{-1}s^{-1}$, $k_{off}$=3.9 x $10^{-1}$ $s^{-1}$) and CTC-445.2d SARS-CoV-1 S RBD ($K_d$=1.2 μM, $k_{on}$=3.7 x $10^5$ $M^{-1}s^{-1}$, $k_{off}$=4.7 x $10^{-1}$ $s^{-1}$). The inset shows the steady state plot.

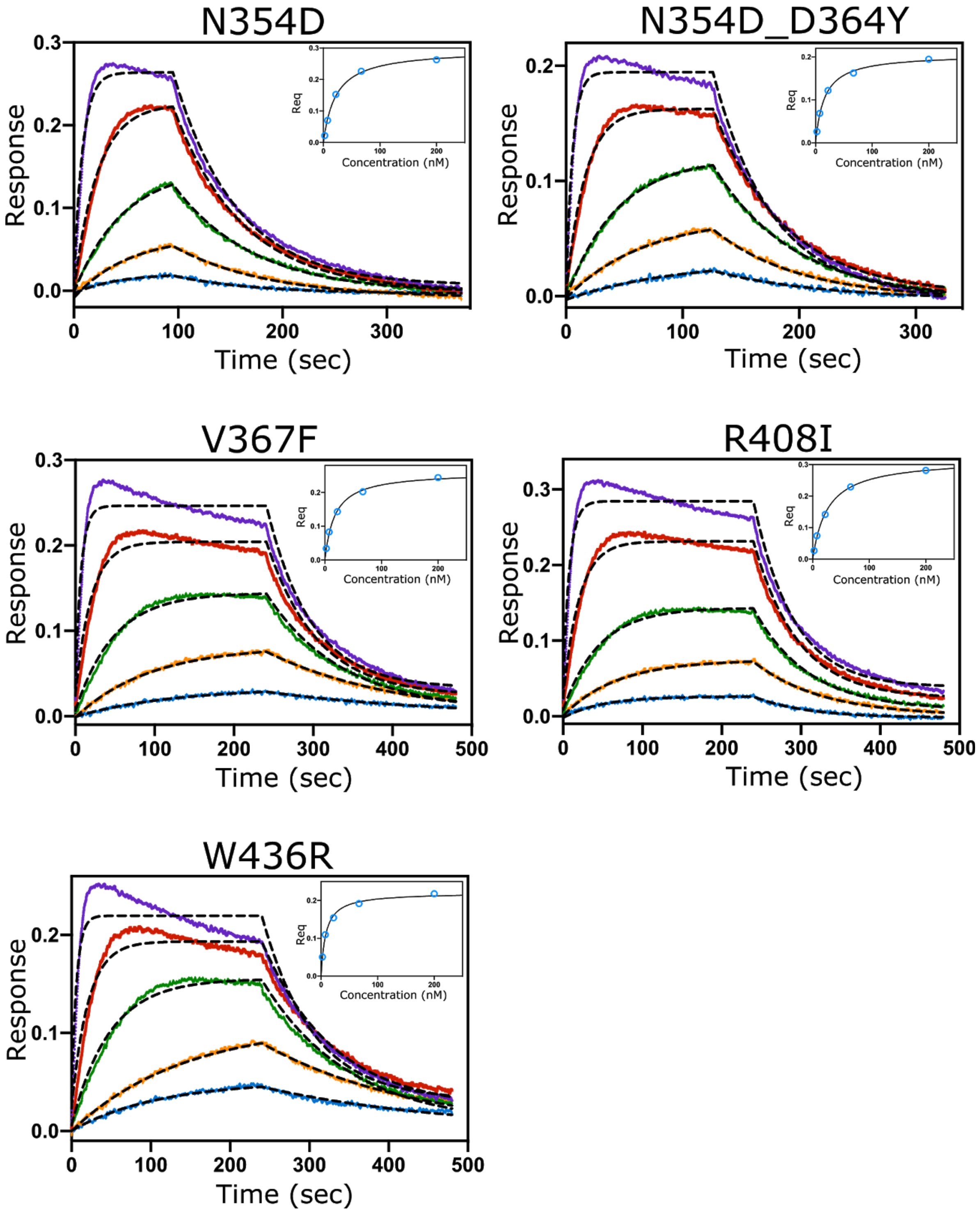


**Figure S10: Kinetics of binding for CTC-445.2 to SARS-CoV-2 RBD mutants.** Five RBD mutants were immobilized via Anti-Penta-His sensors, and CTC-445.2 was titrated in solution. Data was globally fit to a 1:1 model for each RBD mutant. The inset shows the steady state plot.

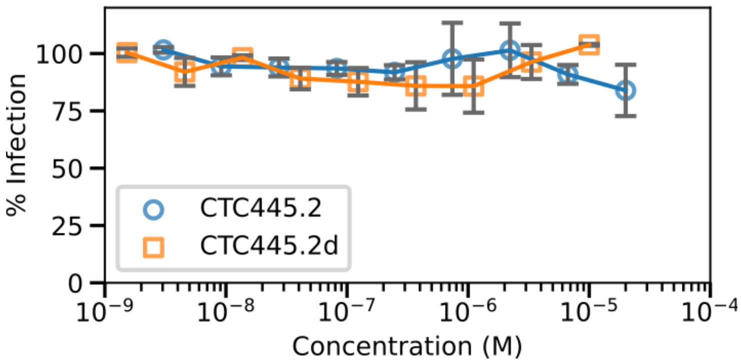


**Figure S11: Inhibition of infection VSV-Luc pseudovirus expressing VSVg by CTC-445.2 and CTC-445.2d.** We observed no inhibition by the de novo decoys, confirming that, as they were designed to, their activity is specific for the SARS-CoV-2 virus.

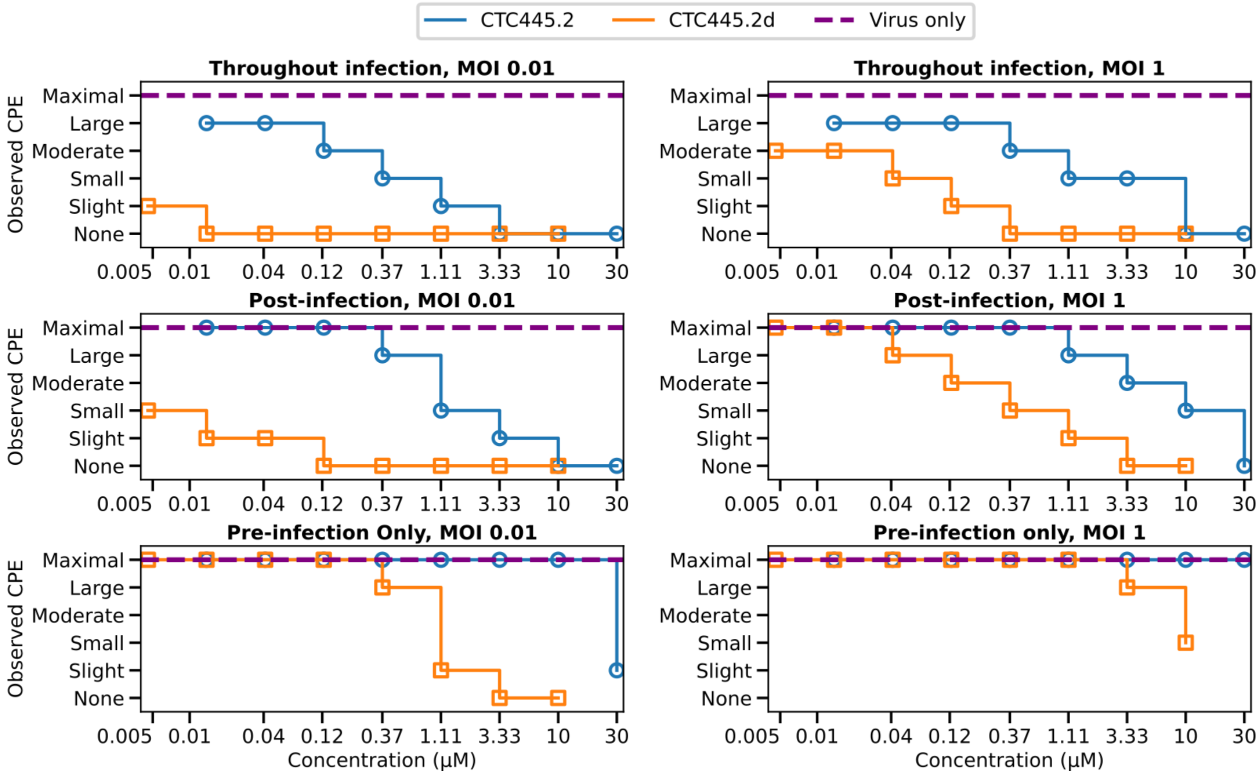


**Figure S12. Inhibition of SARS-CoV-2 (cytopathic effect) in Vero E6 cells.** Observed qualitative cytopathic effect of SARS-CoV-2 infection of Vero E6 cells at 72 hours post infection, where CTC-445.2 or CTC-445.2d is present throughout infection, post-infection only, and pre-infection only.

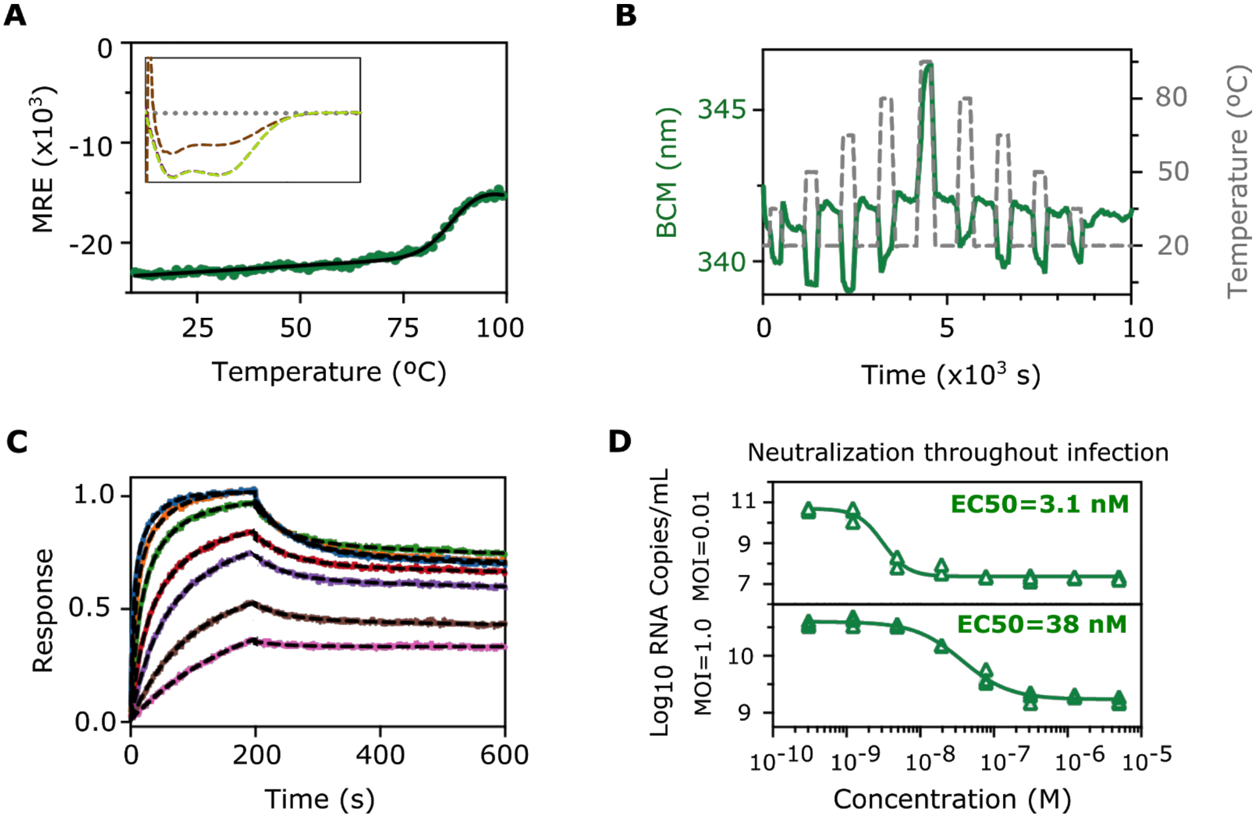


**Figure S13. CTC-445.3d, a further optimized CTC-445.2d variant. Stability, thermal kinetics, binding and in vitro neutralization.** CTC-445.3d is a further stability-enhanced variant. Similarly to CTC-445.2d, CTC-445.3d is a bivalent molecule that contains three additional (i.e. compared to CTC-445.2) amino acid substitutions (A2V, I41L and R142L) in each of the (two) functional domains. **A)** Thermal denaturation of CTC445.3d was followed by circular dichroism at 208 nm, from 20 °C to 99 °C at 2°C/min. A two-state fit revealed a Tm of 89 ± 2 °C, which is 17 °C higher than the Tm of CTC-445.2d. The inset shows the far UV spectra at 20 °C (purple), 99 °C (brown) and 20 °C (green), to show recovery after heating and cooling. **B)** Heating cycles (20, 35, 50, 65, 80, 95 °C, and vice versa down) revealed that CTC-445.3d only loses its secondary structure at the highest heating cycle (95 °C) and goes back to similar BCM values after each temperature cycle. **C)** Binding kinetics of CTC-445.3d to immobilized SARS-CoV-2 RBD (300 nM to 4.7 nM) using octet. Results were fit using a 2:1 binding model to reveal a $K_d$= 1 nM, $k_{on}$= 4.2 x $10^5$ $M^{-1}$ $s^{-1}$, and $k_{off}$= 1.4x $10^{-2}$ $s^{-1}$. **D)** In vitro neutralization of the live BetaCoV/HongKong/VM20001061/2020 SARS-CoV-2 virus by CTC-445.3d in Vero E6 cells, with the protein present throughout the infection and using two MOIs (0.01 and 1.0), three replicates each. Determination of the SARS-CoV-2 RNA copy numbers revealed an EC50 of 3.1 nM at MOI 0.01 and an EC50 of 38 nM at MOI 1.0.

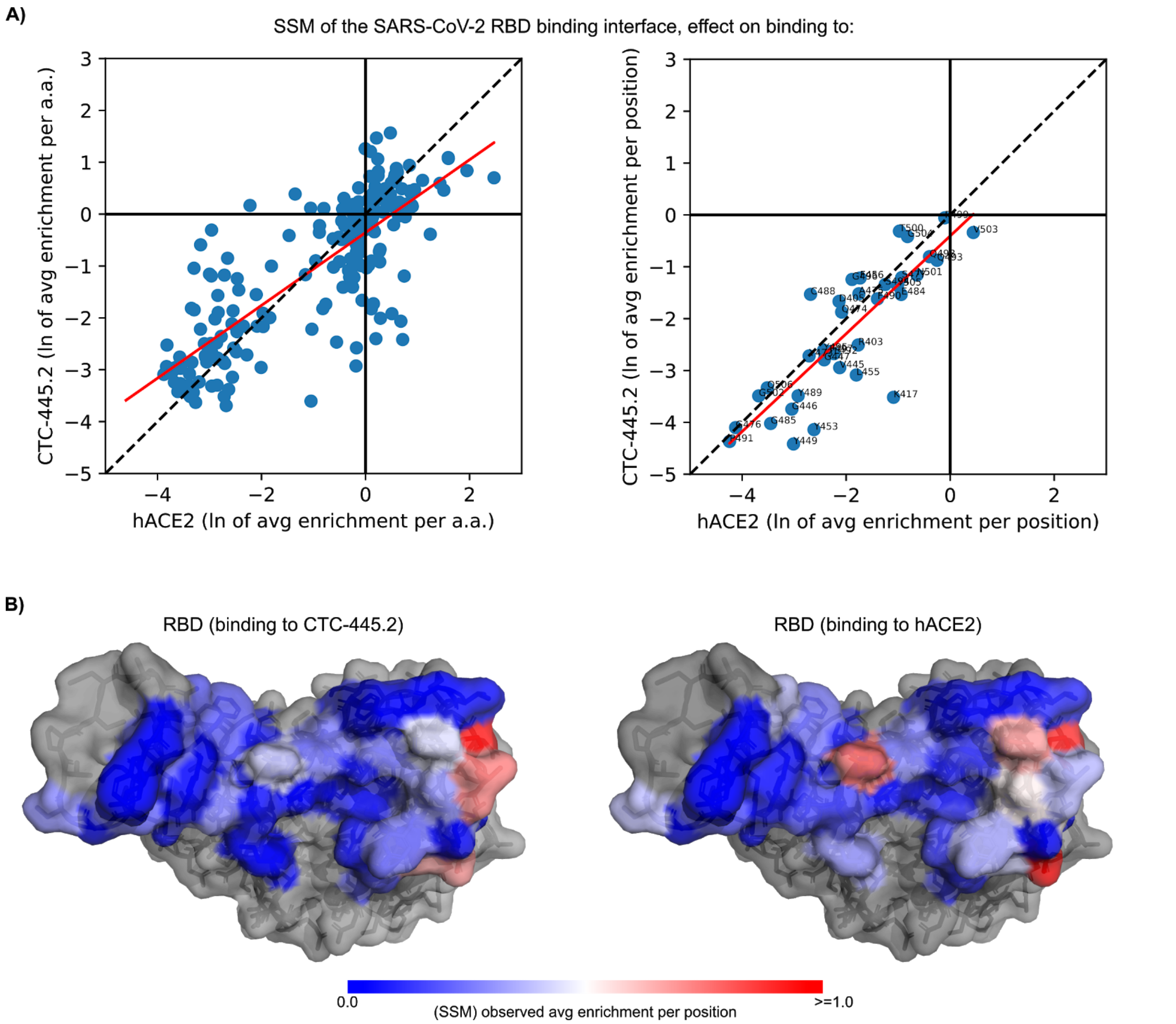


**Figure S14. Yeast-display single-site saturation mutagenesis (SSM) library of the SARS-CoV-2 RBD binding interface, effect on binding of CTC-445.2 and hACE2.** A) The left plot shows the correlation (R2=0.63, red line) of the average effect in enrichment comparing binding by CTC-445.2 *vs.* ACE2. Each data point is a single amino acid mutation at a given position of the RBD. Yeast display FACS for binding was performed at concentrations of: 656.0, 218.0, 72.0, 24.0, 8.0, 2.0, 0.9, 0.3, 0.1 nM of CTC-445.2 or hACE2 (see Methods). Similarly, the right plot shows the correlation (R2=0.71, red line) for the (average) effect per position (i.e. the accumulated effect of any given mutation at a single position in the protein). The dotted line is the trace of a theoretical perfect correlation; B) Shows the mapping of the average effect per position into a surface representation of the RBD, for the binding to (left) CTC-445, and (right) hACE2. As expected for a true de novo decoy, there is a good correlation of the effect of RBD mutations on the binding to both proteins. The surface colored in blue indicates that the overall effect of a mutation in that position is to abrogate binding, while red color indicates the opposite trend. The SSM library was designed to include all the 19 possible natural-amino acid mutations for the 38 residues composing the RBD binding interface with hACE2 (within 10 Å of any Cα or C$\beta$: R403, D405, K417, V445, G446, G447, Y449, Y453, L455, F456, Y473, Q474, A475, G476, S477, E484, G485, F486, N487, C488, Y489, F490, P491, L492, Q493, S494, Y495, G496, F497, Q498, P499, T500, N501, G502, V503, G504, Y505, Q506. About 50% of all possible mutations in the unselected yeast population had sufficient representation to be analyzed. Remarkably ~40% of the mutations exhibited detectable binding to both CTC-445.2 and hACE-2 at the highest concentration tested (656nM).

### Materials and Methods

**Computational *de novo* protein design of ACE2 decoys.** Spike protein RBD-binding motifs interacting with ACE2 were identified and extracted from publicly available structures (PDBs: 6CS2, 6VW1, 6M17); one (“H1”) spanning residues 19-53; another (“H2”) spanning residues 55-84; and in some designs, a third beta-hairpin motif (“EE3”) spanning residues 346-360. The surface-exposed residues from these motifs were preserved, and the buried residues were allowed to change. The supporting structures were either computationally generated and placed by an available method (e.g. Rosetta combinatorial fragment assembly, parametric generation, etc) or extracted from existing structures. The position and orientation of the new elements was refined using a monte carlo search to optimize core packing. This protocol which idealized the motifs and generated fully-connected backbones using a clustered database of known protein structure fragments. A total of 34,589 designs were generated and filtered based on computational metrics (figure 1f). Of the designs that passed the filters, biased forward folding simulations (see below) were run on 1,899 designs to eliminate designs to study the global energy minima (figure 1g). Of the designs that passed the filters, 196 were ordered from four the sets of top-ranked designs in: 1) Rosetta score per residue; 2) biased folding simulations; 3) ddG as computed using Rosetta. In addition, a set of hand-selected designs were synthesized and screened. First, the core elements (i.e., the starting motifs and supporting secondary structure) were rebuilt by identifying parametric equations of repetitive phi and psi angles (omega fixed to 180°) that result in secondary structures that recapitulated each of the target helices as close as possible, a “pitch” on the phi and psi angles was allowed every 3rd residue in order to allow the helices the possibility to have curvature. By using these parametric equations, the algorithm can vary the length of each of the core-elements up to +/- 12 amino acids (compared to the input structural motifs). All length variations of the core-elements were then reconnected pairwise with loops from a clustered database of highly ideal loops (fragment-size of 7 amino acids). To connect pairs of secondary structure elements, the mimetic building protocol aims to reconnect the idealized elements by pairs in all possible combinations. For each pair of secondary structure elements, the loop database was filtered to remove.

The databases of highly ideal fragments used for the design of the backbones for the de novo mimetics were constructed with the Rosetta application “kcenters_clustering_of_fragments” using an extensive database of non-redundant (publicly available) protein structures from the RCSB protein data bank, which was comprised of 7062 PDBs for the 7-mer database.

Folding simulations were biased towards the designed conformation (*44*) by using a small subset of fragments at each residue position with the lowest RMSD (9- and 3-mers) to the designed structure. The funnel-shaped energy landscape suggests that the designed structure is the global energy minima and has a substantial energy gap with respect to alternative conformations. In the plots shown for the simulations, each dot represents an independent biased forward folding trajectory which was computed by monte carlo insertion of fragments from solved protein structures. Three sets of trajectories are computed; those using only the 5 fragments with lowest RMSD to the design (brown), those using fragments from the design model plus the 8 fragments with lowest RMSD (orange) and relax trajectories (blue).

**Yeast display.** Genes encoding *de novo* ACE2 decoys and hACE2 were ordered as gBlocks (IDT), with 60 bases of overlap on each end to the pETcon3 yeast display plasmid. gBlocks and linearized pETcon3 (digested with NdeI and XhoI restriction enzymes and purified by gel extraction) were transformed together into *S. cerevisiae* strain EBY100 cells. Via homologous recombination, the genes for the designed proteins were placed in frame between the *AGA2* gene and the Myc tag on the plasmid, to allow protein expression on the surface of yeast with a c-terminal Myc tag. Yeast were first grown in C-Trp-Ura selective media to saturation, and later induced for 12-18 h in SGCAA media. Cells were then first washed with chilled PBSF buffer (50 mM $NaPO_4$, 150 mM NaCl, 0.1% w/v BSA, pH 7.4), and allowed to bind to 200 nM SARS-CoV-2 Spike RBD-mFc (Sino Biological; residues Arg319-Phe541), for 30 min on ice. Cells were washed with chilled PBSF buffer again, and tagged with FITC-conjugated chicken anti-cMyc antibody (ICL), at 0.8 µL per $10^6$ cells, and phycoerythrin-conjugated goat anti-mFc antibody (Jackson ImmunoResearch), at 1.6 µL per $10^6$ cells, for 15 min on ice, and washed again with chilled PBSF buffer. Lastly, cells were analyzed by flow cytometry using the Guava easyCyte flow cytometer.

To characterize the binding affinity and competition to hACE2 of CTC-445, yeast display assays were performed as described above, using varying amounts of SARS-CoV-2 Spike RBD-mFc (0-243 nM) for the titration assay and 9 nM SARS-CoV-2 Spike RBD-mFc with varying concentrations of soluble human ACE2 His-tag (Sino Biological; residues Met1-Ser740) (0-729 nM) for the competition assay.

**Directed evolution.** Error prone PCRs (epPCR) of the gene coding for CTC-445 were performed using a GeneMorph II Random Mutagenesis Kit (Agilent), according to the standard protocol. Two separate PCR reactions were performed using 10 ng and 1 ng of the CTC-445-coding gene as the initial target DNA, in order to obtain a higher and lower error rate pool. Amplification was performed with the pETCON_overlap_fwd (AGTGGTGGAGGAGGCTCTGGTGGAGGCGGTAGCGGAGGCGGAGGGTCGGCTAG CCATATG) and pETCON_overlap_rev (AGATCTCTATTACAAGTCCTCTTCAGAAATAAGCTTTTGTTCGGATCCGCCCCCCT CGAG) primers. Thirty cycles of amplification were used for the PCR, with an annealing temperature of 55°C. The products of both PCRs were combined, and purified by ethanol precipitation. 12 µg of CTC-445 epPCR DNA, together with 4 µg of linearized pETcon3 vector, were transformed by electroporation into conditioned *S. cerevisiae* strain EBY100 cells. The transformation efficiency was $2x10^6$ transformants.

Six rounds of cell sorting were performed on the library of CTC-445 epPCR, to identify mutations that improve binding to SARS-CoV-2 and improve its stability to proteases. Yeast were grown in C-Trp-Ura selective media and later induced for 12-18 h in SGCAA media. Cells were pretreated before primary labeling with a mixture of trypsin (12.5 µg/mL) and chymotrypsin (5 µg/mL) in TBS buffer (25 mM Tris-HCl, 150 mM NaCl, pH 8.0) for 5 min at room temperature. This reaction was halted by adding a large excess volume of ice-cold TBSF 1% (25 mM Tris-HCl, 150 mM NaCl, 1% w/v BSA, pH 8.0) and washing the cells four additional times with TBSF 1%. Cells were then labelled with decreasing concentrations of SARS-CoV-2 Spike RBD-mFc at every round of sorting: 200 nM (round 1, $10^8$ cells sorted), 10 nM (round 2, $10^7$ cells sorted), 1 nM (round 3, $10^6$

cells sorted), 100 pM (round 4, $10^6$ cells sorted), 10 pM (round 5, $10^6$ cells sorted) and 1 pM (round 6, $10^6$ cells sorted), for 30 min on ice, and washed with chilled PBSF buffer. Starting at sorting round 4, an additional selection step for improved $k_{off}$ was added, where cells were incubated in 1 ml PBSF buffer at 37 °C for 1 hour, and washed again with chilled PBSF buffer. Lastly, cells were incubated with FITC-conjugated chicken anti-cMyc antibody (ICL), at 0.3 µL per $10^6$ cells for sorts 1 and 2, and 3 µL per $10^6$ cells for remaining sorts and phycoerythrin-conjugated goat anti-mFc antibody (Jackson ImmunoResearch), at 0.4 µL per $10^6$ cells for sorts 1 and 2, and 4 µL per $10^6$ cells for remaining sorts, for 15 min on ice, and washed again with chilled PBSF buffer. Fluorescence-activated cell sorting (FACS) was performed with the Sony SH800 instrument, selecting the top 5% of the displaying subpopulation by their PE/FITC fluorescence ratios. Lastly, selected cells were plated on SDCAA agar plates and single colonies were picked, amplified by PCR, and sequenced by Sanger sequencing (Genewiz).

**Cloning and molecular biology.** The dsDNA CTC-445 gBlock with pETcon3 overlaps (5' overlap: 5'-gctagtggtggaggaggctctggtggaggcggtagcggaggcggagggtcggctagccatatg-3', 3' overlap: 5'-ctcgagggggggcggatccgaacaaaagcttatttctgaagaggacttgtaatagagatct-3') was digested with NdeI and XhoI endonucleases (New England Biolabs) for 1 hour at 37°C followed by ligation into linear pET29b(+) using T4 DNA Ligase (New England Biolabs). The vector was linearized by 100-fold overdigestion by NdeI and XhoI and then purified by gel extraction (Qiagen). For the optimized CTC-445 variants, genes with overlaps for pET29b(+) (5': AAATAATTTTGTTTAACTTTAAGAAGGAGATATACATATG, 3': CTCGAGCACCACCACCACCACCACTGAGATCCGGCTGCTA) were designed, ordered and cloned by overlap extension or Gibson assembly (*45*). To facilitate protein quantification, and to enable cleavage of the C-terminal His tag, a C-terminal GSGWGSGLVPRGS sequence was added to all CTC-445-derived sequences.

**Recombinant protein expression.** Protein sequences were synthesized (IDT) and cloned into pET-29b(+) E. coli plasmid expression vectors (both with and without a thrombin-cleavable C-terminal 6- His tag). Plasmids containing the clones of CTC-445, one non-expressing negative control and a positive control protein transformed into chemically competent E. coli Lemo21 cells (NEB). Protein expression was then induced with 1 mM of isopropyl β-d-thiogalactopyranoside (IPTG) at 37 °C. After 4 h expression, samples were collected and run on SDS-PAGE gel along with a PageRuler Plus Prestained Protein Ladder.

**Chemical protein denaturation.** Samples of CTC-445, CTC-445.2 and CTC-445.2d were incubated overnight at 1 mg/mL with 0-6 M guanidinium chloride in PBS buffer (pH 7.4). Circular dichroism spectra were measured for each sample at 25 °C in a 0.1 mm path-length cuvette using an Chirascan spectropolarimeter V100 (Applied Photophysics). The unfolding curve at 222 nm was fitted to the following equation:

$$fD = \frac{\left((yN+mN[GuHCl])+(yD+mD[GuHCl])*exp^{-\left(\frac{\Delta GND-mND[GuHCl]}{RT}\right)}+(yI+mI[GuHCl])*exp^{-\left(\frac{GID-mID[GuHCl]}{RT}\right)}\right)}{\left(1+exp^{-\left(\frac{\Delta GND-mND[GuHCl]}{RT}\right)}+exp^{-\left(\frac{\Delta GID-mID[GuHCl]}{RT}\right)}\right)}$$

Where fD is the fraction of protein in the unfolded state, yN, yI and yD are the signals of the native, intermediate and unfolding states, mN, mI and mD represent the signal dependence with the denaturant concentration, ΔGND and ΔGID are the free energy changes for the native and intermediate states while mND and mID represent their dependence with de denaturant concentration. Finally, R represents the gas constant in kcal $mol^{-1}$ $K^{-1}$ and T the temperature in K. ΔGNI is calculated from the subtraction of ΔGID to ΔGND.

**Circular dichroism.** Far-ultraviolet circular dichroism measurements were carried out with an CHIRASCAN spectrometer V100 (Applied Photophysics) in PBS buffer (pH 7.4) in a 0.1 mm path-length cuvette with protein concentration of 0.2 mg/mL (unless otherwise mentioned in the text). Temperature melts were obtained from 20 to 95 °C and monitored absorption signal at 222 nm (steps of 0.5 °C per min, 30 s of equilibration by step). Wavelength scans (195–250 nm) were collected at 20 °C, 95 °C, and again at 20 °C after refolding (roughly 5 min). Melting temperature (Tm) values were calculated from the fitting of the thermal melts to the following equation:

$$fD = \frac{\left((yN+mNT)+(yD+mDT)*exp\left[\frac{\Delta HvH}{R}\left(\frac{1}{T}-\frac{1}{Tm}\right)\right]\right)}{\left(1+exp\left[\frac{\Delta HvH}{R}\left(\frac{1}{T}-\frac{1}{Tm}\right)\right]\right)}$$

Where fD is the fraction of the protein in the unfolded state, HvH is the enthalpy change associated to the unfolding process, yN and mN represent the signal of the native state and its dependence with the temperature, yD and mD represent the signal of the unfolded state and is dependence with the temperature and R is the gas constant.

**Serial thermal ramping protein stability.** Serial thermal ramping assays were performed in order to test the unfolding reversibility (thermal recovery) of CTC-445, CTC-445.2, CTC-445.2d and hACE2 (Sino Biological); to this end, the UNCLE platform (Unchained Labs) was used. 8.8 µL of CTC-445, CTC-445.2 and CTC-445.2d at 0.5 mg/mL, and hACE2 at 0.25 mg/mL, were loaded in capillary cuvettes and fluorescence spectra (~300-400 nm) were measured for each sample while temperature was increased and decreased repeatedly. Temperature ramps were performed in the following order: 20 °C to 35 °C, 20 °C to 50 °C, 20 °C to 65°C, 20 °C to 80 °C, 20 °C to 95 °C, 20 °C to 80 °C, 20 °C to 65 °C, 20 °C to 50 °C and 20 °C to 35 °C. All samples were in PBS buffer pH 7.4. The barycentric mean was calculated for each fluorescence spectrum using the UNCLE analysis program (Unchained Labs).

**Size-exclusion chromatography with multi-angle light scattering (SEC-MALS)**. SEC-MALS assays were performed using a 1260 infinity II LC HPLC system (Agilent Technologies) with a Superdex 75 10/300 size exclusion column (GE Healthcare), coupled to an inline static light scattering instrument (MiniDawn, Wyatt Technology) and differential refractive index (Optilab rEX, Wyatt Technology) and UV detection systems. Protein samples were prepared in PBS buffer at concentrations ranging from 2 to 4 mg/mL and filtered with 0.22 µm syringe filters; 100 µL of the samples were injected and ran at a flow rate of 0.5 mL/min. Data were analyzed using the software ASTRA 7 (Wyatt Technologies).

**ELISA competition assays.** SARS-CoV2 S Protein RBD (Acro, Cat#EP-105) was coated at 0.5 µg/mL (100ul/well) in flat, clear bottom, high binding polystyrene 96-well plate (Thermo, Cat#15041) overnight at 4°C. On the following day, each assay plate was washed 3X with 0.05% Tween-PBS and blocked with 2% BSA/0.05%Tween-PBS for 1-1.5 hours at 37°C. On the following day, 11-point four-fold serial dilutions of each test article were prepared starting at 2 µM (2X final). In addition, a dose response of biotin-hACE2 was prepared by performing a two-fold serial dilution starting at 0.174 µg/mL (2 nM). From the 0.174 µg/mL biotin-hACE2, a constant concentration of 0.07 µg/mL (0.8 nM, 2X final) was prepared to be used for inhibition by test articles. After blocking, each assay plate was washed (3X) and each test article dilution was added to the plate at 50 µL/well (single replicate well per concentration) followed by 50 µL/well of biotin-hACE2 (~0.035 µg/ml, equivalent to a concentration of 0.4 nM). Incubation of test articles with 0.4 nM biotin-hACE2 was performed for 1 hour at 37°C. Afterwards, each assay plate was washed (3X), followed by incubation of 0.1 µg/ml (100 µL/well) streptavidin-HRP for 1 hour at 37 °C. Lastly, each plate was washed (3X) before addition of 100 µl/well TMB (abCam, Cat#171524). TMB was incubated for 7-8 minutes and the reaction was stopped with the addition of 50 µL/well TMB Stop Solution (abCam, Cat# ab171529). Mean absorbance in each well was measured from 4 spots at 450 nm.

**Biolayer Interferometry.** Binding function of CTC-445 was analyzed by biolayer interferometry. Binding data were collected in an Octet RED96 (ForteBio) and processed using the instrument's integrated software using a 1:1 binding model. SARS-CoV-2 Spike Protein (RBD domain only, his Tag, Sino Biological) were immobilized to Anti-Penta-His sensors (HIS1K, ForteBio) at 2 µg/mL in binding buffer (10 mM HEPES, pH 7.4, 150 mM NaCl, 3 mM EDTA, 0.05% surfactant P20, 0.5% non-fat dry milk). After baseline measurement in the binding buffer alone, the binding kinetics were monitored by dipping the biosensors in wells containing defined concentrations of the designed protein (2.4-200 nM) (association) and then dipping the sensors back into baseline wells (dissociation).

**Deep mutational scanning / Single-site saturation mutagenesis (SSM).** Positions in the SARS-CoV-2 S protein RBD within 10Å of the ACE2 interface were mutated by PCR mutagenesis using custom primers (IDT, Coralville, IA) to introduce an NNK codon at each position of interest, as described previously (*46*). PCR products were purified using a QiaQuick gel extraction kit (Qiagen, Hilden, Germany), and 4 µg of the resulting site-saturation mutagenesis library was transformed along with 1 µg of linearized pETcon3 vector into EBY100 yeast via electroporation with a final transformation efficiency of $1x10^6$ transformants, as described previously (*47*). The resulting library was expressed by surface display and sorted by FACS at varying concentrations of CTC-445.2 or hACE2 (656.0, 218.0, 72.0, 24.0, 8.0, 2.0, 0.9, 0.3, 0.1 nM), as described for the directed evolution experiment, with all cells showing binding fluorescence above baseline collected for downstream analysis. In addition, a nonbinding pool was collected representing all cells that showed expression in the absence of either CTC-445.2 or hACE2. All sorted libraries, as well as an unselected library were processed to recover plasmid DNA, and and was then amplified via two rounds of PCR followed by AMPure bead cleanup (Beckman Coulter, Indianapolis, IA) using custom primers to add Illumina sequencing primer bind sites, barcodes, and flow cell adapters, as previously described (*48*). Prepared libraries were then sent for Illumina sequencing (GENEWIZ, South Plainfield, NJ), with a yield of between ~895k and ~3248k reads per library. Enrichment was evaluated for mutations at the following positions in the SARS-CoV-2 S protein RBD: 403, 405, 417, 445, 446, 447, 449, 453, 455, 456, 473, 474, 475, 476, 477, 484, 485, 486, 487, 488, 489, 490, 491, 492, 493, 494, 495, 496, 497, 498, 499, 500, 501, 502, 503, 504, 505, and 506. Sequencing data was aligned to the DNA sequence for the RBD using the bowtie2 aligner (33 bases were trimmed from the 5' end and 30 from the 3' end); any aligned sequences containing insertions or deletions were discarded. Sequencing data was filtered to remove bases with phred quality score $Q<15$. To ensure sufficient coverage to compute enrichment, mutations with less than 20 counts in the expression-only dataset were removed. To determine enrichment, the number of occurrences in each dataset of each possible position and amino acid were first computed. To compute the fraction of reads each amino acid and position were present in each dataset in a way that is robust to outliers, the number of occurrences data was scaled to the interquartile range between the 1st quartile (25th quantile) and the 3rd quartile (75th quantile) using the RobustScaler from the scikit-learn python module. Next, enrichment factors for all amino acid substitutions were determined. For each dataset, enrichment was calculated as the scaled counts for each dataset divided by the scaled counts for the expression-only dataset, giving 9 enrichment values for each position and amino acid corresponding to the 9 concentrations tested for both CTC-445.2 and ACE2. These 9 enrichment values were smoothed by averaging the lowest 3, middle 3, and highest 3 concentrations. The natural logarithm of the maximum enrichment of these smoothed values for ACE2 and CTC-445.2 was used as an estimate of the effect of each amino acid substitution on binding. Amino acids and positions with 0 counts in each dataset were removed.

**VSV-luc pseudovirus neutralization.** Neutralization activity was determined using a non-replicative VSV pseudovirus containing a firefly luciferase gene and expressing the spike protein from Wuhan-1 SARS-CoV-2 isolate (GenBank: QHD43416.1).

Neutralization assays were performed with either SARS-CoV-2 / VSV-Luc (expressing the full ectodomain of the SARS-CoV-2 spike protein on the surface of the pseudovirus) or VSVg / VSV-Luc (expressing VSVg). Thirty thousand 293T-ACE2 cells were seeded the day before in white plates in the presence of hygromycin (100 µg/mL). The day of the neutralization assay, 25 µL of media containing pseudovirus was mixed with 25 µL of serial dilutions of the test-item in a different plate, and then incubated for 1 h at 37°C. Pseudovirus and test-item dilutions were performed in DMEM media containing 5% heat-inactivated fetal bovine serum (DMEM5). After the 60 minute incubation, the test-item / pseudovirus mixture was added to the 293T-ACE2 cells. Infection was allowed for twenty-four hours. Firefly luciferase activity was monitored at 24h using the Britelite reporter gene assay (Perkin Elmer).

Test-items were evaluated in duplicate using serial 3-fold dilutions. Controls included cells infected with a VSV pseudovirus lacking spike protein (NoEnv / VSV-Luc), with pseudovirus carrying VSVg (VSVg / VSV-Luc), or uninfected cells ("mock-infected"). Pseudovirus-infected cells were incubated with test-item or vehicle alone (DMEM5). Some wells included cells challenged with pseudoviruses incubated with dilutions of plasma from a convalescent patient recovered from COVID-19 (COV+) or with a control plasma (COV-). Before collecting blood from COV+, this patient had been confirmed infected with SARS-CoV-2 by RT-PCR, and also for the presence of IgGs against SARS-CoV-2 Spike (S) evaluated with a lateral flow antibody test. COV- plasma was derived from a PCR-negative and IgG-negative for S antibodies.

Average RLU values in mock-infected cells were subtracted from all values ("mock-subtracted values"). The average RLU value of infected samples in the presence of vehicle was obtained. For each data point, once the mock was subtracted, the value was divided by the average value in infected cells in the presence of vehicle alone and then multiplied by 100 to obtain the percentage infectivity value normalized to samples with vehicle alone. The average percentage infectivity for each concentration together with the standard deviation of duplicates was obtained from duplicate values generated as shown in point 4.

The 50% inhibitory concentration (IC50) was determined with GraphPad Prism. The IC50 was defined as the concentration of test-item at which the relative luminescence units (RLUs) were reduced by 50% as compared with the values in cells infected with pseudovirus in the absence of test-item).

**Viral infection assay of the SARS-CoV-2 virus in Vero E6 cells.** Vero E6 cells (ATCC #CRL-1586) were maintained in Dulbecco's Modified Eagle Medium (DMEM) supplemented with 4.5 g/L D-glucose, 100 mg/L sodium pyruvate, 10% FBS, 100,000 U/L penicillin–streptomycin and 25 mM HEPES. SARS-CoV-2 virus, BetaCoV/Hong Kong/VM20001061/2020, was isolated from a confirmed COVID-19 patient at the end of January 2020 and was passaged 3 times in Vero E6 cells. All experiments using the SARS-CoV-2 isolate were performed at the core BSL3 facility, LKS Faculty of Medicine, The University of Hong Kong. The inhibitory effects of CTC-445.2 and CTC-445.2d were firstly evaluated with SARS-CoV-2 at the multiplicity of infection (MOI) of 1.0 or 0.01 TCID50/cell in 96-well plates. Three-fold serially diluted CTC445.2 (final concentrations ranged from 30 µM to 0.01 µM) and CTC445.2d (final concentrations range from 10 µM to 0.005 µM) were pre-incubated with diluted SARS-CoV-2 virus for 1 hour at 37°C and the mixture was added to Vero E6 and further incubated 1 hour at 37°C. The inoculum was removed and cells were overlaid with fresh infection media (DMEM supplemented with 4.5 g/L D-glucose, 100 mg/L sodium pyruvate, 2% FBS, 100,000 U/L Penicillin-Streptomycin, and 25 mM HEPES) containing matched concentrations of CTC-445.2 and CTC-445.2d. After 72 hours incubation at 37°C, culture supernatants were collected to determine viral load (viral RNA copies/mL) using TaqMan™ Fast Virus 1-Step Master Mix as described (*49*). Four-parameter logistic regression (GraphPad Prism) was used to fit the dose-response curves and determined $EC_{50}$ of CTC-445.2 and CTC-445.2d. To confirm the mechanism of inhibition, time-of-addition assay was used to evaluate the inhibitory effect of CTC-445.2 and CTC-445.2d. To test the effect of the, decoy proteins in blocking SARS-CoV-2 virus entry, 3-fold serially diluted CTC-445.2 and CTC-445.2d were pre-incubated with SARS-CoV-2 for 1 hour at 37°C and the mixture was added to Vero E6 cells and further incubated for 1 hour at 37°C. The inoculum was then removed and the cells were washed in PBS three times before fresh infection media (without CTC-445.2 or CTC-445.2d) was added to the cells. To test the effect of the decoy protein in blocking SARS-CoV-2 under multi-cycle replication, SARS-CoV-2 virus was allowed to incubate with Vero E6 cells for 1 hour at 37°C; the inoculum was removed and the cells were washed in PBS three times before infection media containing 3-fold serially diluted CTC-445.2 and CTC-445.2d were added. Culture supernatants were collected at 72 hours post-infection to determine viral load (viral RNA copies/mL). Dose-response data was fit to a three-parameter nonlinear fit using the scipy python module (*50*), with the upper bound of the curve fixed to the experimentally determined data points at concentration=0 μM.

**Viral infection assay of the (NanoLuc) icSARS-CoV-2-nLuc virus in Calu-3 cells.** Calu-3 (ATCC HTB-55) from American Type Culture Collection (ATCC, Manassas, VA) was maintained in growth medium comprising OptiMEM with 20 mM HEPES, 10 mM GlutaMAX, 4% FBS, 10,000 IU/mL Penicillin, 10,000 μg/mL Streptomycin, and 10ng/mL Epidermal Growth Factor (EGF). Vero cells (USAMRIID) and icSARS-CoV-2-nLuc) (Isolate USA-WA1/2020) were generously provided by Dr. Ralph Baric Lab at University of North Carolina at Chapel Hill (Hou et al., Cell. 182, 1-18. 2020). icSARS-CoV-2-nLuc virus stock was propagated in Vero cells and stored at -80 °C. Calu3 cells are seeded in white 96-well flat-bottomed tissue culture plates. All viral infections were performed in the biosafety level 3 facility at the University of Washington. Three-fold serial dilution of test compound CTC-445.2 (final concentration ranging from 20 μM to 0.01 μM) and CTC-445.2d (final concentration ranging from 10 μM to 0.0046 μM) were added to corresponding wells during virus infection at MOI of 1.0; three independent experiments were performed in duplicate. After virus adsorption at 37℃ for 1 h, inoculum was removed and replaced with fresh growth media containing the same compound concentration to maintain the treatment throughout the entire infection course. To estimate the antiviral effects of test compounds, icSARS-CoV-2-nLuc virus replication was measured by adding Nano-Glo Endurazine

Live Cell substrate to each well, and nanoLuc readings of the same well were taken at 24, 48, and 72 hours post-infection. NanoLuc reads were normalized to the untreated wells, and the inhibitory effects of the compounds on virus replication were analyzed by non-linear regression curve fit algorithm by using GraphPad Prism.

**Cytotoxicity assay**

Cytotoxicity of the test compounds in treated Vero-E6 or Calu-3 cells was tested using the Cell Counting Kit-8 (CCK-8) assay (Abcam). Cells were seeded in 96-well flat-bottom tissue culture treated plates. Four-fold serial dilutions of test compound CTC-445.2 and CTC-445.2d in fresh infection medium (final concentrations ranging from 60 uM to 3.7 nM for CTC-445.2 and 50 μM to 3nM for CTC-445.2d, respectively) were added onto corresponding Vero E6 cells. Three-fold serial dilutions of the test compounds of CTC-445.2 (final concentration ranging from 20 μM to 0.01 μM) or CTC-445.2d (final concentration ranging from 10 uM to 0.0046 uM) in Calu-3 cell growth media were added onto the cells. Cells were then incubated at 37℃, 5% $CO_2$ for 2 days. 10 μL CCK-8 reagent was added to each well, and the absorbance was measured at 460 nm.

**Pharmacokinetics of the de novo decoys.** Eight week old Balb/c mice (Charles River) were anesthetized with isoflurane and 30 μL of CTC-445.2d was delivered intranasally. Mice were euthanized at indicated time points and whole blood and lungs isolated. Whole blood was centrifuged, and plasma was separated from cells by pipetting into a separate tube before freezing. Lung lysate was prepared through mechanical disruption of tissue using Precellys tissue homogenizer (Bertin), followed by lysis with T-PER tissue lysis buffer (ThermoFisher) containing protease/phosphatase inhibitor cocktail (ThermoFisher). Lysates were cleared by centrifugation and were frozen for analysis. A standard 96-well MSD (L15XA) plate was pre-coated with 50 μL/of 0.5 μg/mL SARS-CoV-2 S Protein RBD tagless (Sino, 40592-VNAH) overnight at 4 ℃ and sealed. Dilution of capture reagent was prepared in PBS. On the following day, the assay plate was washed (6X) with 0.05% Tween-PBS and blocked with 150 μL/well MSD Blocker A (5% protein-PBS solution) for a minimum of 2 hours at RT. Plate shaking was performed for all incubation periods. During blocking incubation, a standard curve was prepared by performing a 4-fold serial dilution of CTC-445.2d in either untreated lung lysate or untreated plasma at 10,000 ng/mL. Prior to addition to the assay plate, each sample and standard was diluted 5-fold (MRD = 5) into assay dilution buffer (0.5% MSD Blocker A). In addition, samples were diluted 10-fold in separate wells. After blocking, an assay plate was washed (6X) and 50 μL/well of each sample or standard dilution was added. Assay plate was incubated for additional 1 hour at RT with plate shaking followed by washing (6X). For detection, a 1-step detection method was used. With 1-step detection, rabbit anti-His mAb (RevMab, 31-1048-00)) and SULFOTAG goat anti-rabbit IgG (H+L) pAb (MSD, R32AB) were pre-mixed at 1 μg/mL each for 1 hour prior to addition to assay plate at 50 μL/well. After the last wash, 150 μL/well of MSD Gold Read Buffer was added and luminescence measured on the MSD Quickplex

SQ120. Luminescence was converted to concentration based on a standard curve.

**ACE2 functional assay.** Inhibition of ACE2 enzymatic activity was performed using the ACE2 Inhibitor Screening Kit (Promocell Heidelberg, Germany). 50 µL of ACE2 enzyme solution was added to each well of a 96-well white walled plate (Corning, New York). Test samples were diluted to 20 µM in the provided assay buffer for the highest concentration, with an 8 point 1 to 4 serial dilution. DX600 was used as the ACE2 inhibitor positive control. 10 µL of diluted samples were added to the ACE2 enzyme solution and incubated for 15 min at room temperature. 40 µL of ACE2 substrate mix was then added to each well. After 15 min, fluorescence was measured in an Infinite M1000 plate reader (Tecan Männedorf, Switzerland) with 320 nm excitation and 420 emission filters. Fluorescence values were normalized using negative controls.
